# A network-based integrated framework for predicting virus-host interactions

**DOI:** 10.1101/505768

**Authors:** Weili Wang, Jie Ren, Kujin Tang, Emily Dart, Julio Cesar Ignacio-Espinoza, Jed A. Fuhrman, Jonathan Braun, Fengzhu Sun, Nathan A. Ahlgren

## Abstract

Metagenomic sequencing has greatly enhanced the discovery of viral genomic sequences; however it remains challenging to identify the host(s) of these new viruses. We developed VirHostMatcher-Net, a flexible, network-based, Markov random field framework for predicting virus-host interactions using multiple, integrated features: CRISPR sequences, sequence homology, and alignment-free similarity measures (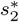 and WIsH). Evaluation of this method on a benchmark set of 1,075 known viruses-host pairs yielded host prediction accuracy of 62% and 85% at the genus and phylum levels, representing 12-27% and 10-18% improvement respectively over previous single-feature prediction approaches. We applied our host-prediction tool to three metagenomic virus datasets: human gut crAss-like phages, marine viruses, and viruses recovered from globally-distributed, diverse habitats. Host predictions were frequently consistent with those of previous studies, but more importantly, this new tool made many more confident predictions than previous tools, up to 6-fold more (n*>*60,000), greatly expanding the diversity of known virus-host interactions.

## Background

Viruses are the most abundant and highly diverse biological entities on earth [1, 2]. Viruses infect all domains of life including archaea, bacteria, and eukaryotes. For prokaryotic viruses, especially those that infect bacteria, there have been extensive studies about their diversity [3, 4], functions [5, 6, 7], and impact on microbial communities through virus-host interactions [8, 9, 10, 11]. In particular, prokaryotic viruses can significantly impact human health [12, 13, 14] and the functioning of many ecosystems [15, 16, 17] such as marine and soil habitats. Therefore, characterizing virus-host interactions is a critical component to understanding how biological systems work. Viruses are traditionally studied using culture-based isolation techniques that provide direct identification of virus-host pairs. Isolation approaches are, however, low throughput and limited to hosts that are cultivable. Compared to the predicted number of extant viruses, a relatively small number of viruses have been discovered via isolation based approaches with current estimates indicating that 75-85% of viruses remain uncharacterized [11, 18]. With the advent of metagenomic sequencing technologies, genetic material from microbes including viruses, regardless of cultivability, can be sequenced. Metagenomic shotgun sequencing, especially the metagenomic sequencing of virus-like particles, has tremendously accelerated the discovery of previously unknown viruses. An example is crAss-like phages, a highly abundant family of ubiquitous human gut viruses, originally discovered from the cross-assembly of fecal viral metagenomic samples [19].

Identifying the hosts of viruses is important for understanding the impact of viruses on the host dynamics and thus host community diversity and function. Computational methods have been developed to infer the hosts of new viruses. Many bacteria and archaea possess CRISPR virus defence systems whereby the host incorporates some virus DNA fragments into its own genome forming interspaced short palindromic repeats (CRISPR) spacers. Therefore, shared CRISPR regions are direct evidence supporting virus-host interactions [16, 19]. Genome alignment matches between virus and host genomes due to integrated prophages or horizontal gene transfer are another piece of strong evidence used in predicting the host of a virus [5, 16]. However, the above methods are limited by their low accessibility: it is estimated that CRISPRs are only present in approximately 10% of sequenced bacterial genomes [20, 21]; many viruses infect hosts under lytic mode without integration to the host genome; and many viruses do not extensively share host genes. Thus, CRISPRs and alignment-based approaches are not applicable for predicting many viral hosts.

Several investigators have utilized the fact that viruses are often more similar to their hosts, compared to non-host species, in terms of genome-wide signature, i.e. *k*-mer usage [11, 22, 23, 24]. This has been used to predict the host of a virus as the one closest to the viral genome based on some similarity measures using *k*-mers. These methods in general have decent prediction accuracy, though the mechanism behind this phenomenon is not fully understood. One plausible explanation is that viruses tend to adopt the codon usage of their hosts in order to utilize the hosts’ translational machinery [25, 26]. The recently developed dissimilarity measure 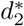 that subtracts expected *k*-mer frequency from the observed frequency achieves the highest reported host prediction accuracy among all current genomic signature-based measures, including commonly used Euclidean and Manhattan distances [23]. Similarly, Galiez *et al.* [24] predicted the host of a virus to be the one for which results from a Markov chain model analysis had the highest likelihood score. The method has good accuracy for short viral fragments. These genomic signature-based measures are often referred to as alignment-free sequence comparison measures. The high correlation between virus and host abundance profiles across different samples also serves as evidence for virus-host interaction [19], but its accuracy is not as high as the above methods [22]. Edwards *et al.* [22] recently provided a comprehensive evaluation of several different computational approaches for virus-host predictions.

In addition to the methods using features defined between a pair of virus and host genomes, some researchers have used virus-virus similarity networks to infer the host of a query virus [27, 28]. The high similarity between viruses may indicate a common host or very close host relatedness. Network-based prediction models, whereby unknown entities are predicted based on the features of their neighbors in a network, have been successfully applied to many biological problems, including predicting protein functions using protein-protein interaction networks [29, 30], inferring disease genes based on gene-gene networks [31, 32, 33], and predicting the target of new drugs using drug-drug, drug-target and target-target similarity networks [34]. A few attempts have been made to exploit the possibility of predicting viral hosts based on virus-virus network information. Different principles, such as gene homology [35, 36], protein family [37] and genome similarity [27, 38, 39] were used to define the virus-virus relationships in networks. Villarroel *et al.* proposed HostPhinder [27], a method to predict the host of a virus by searching for the virus that shares the most *k*-mers from a database of viruses with known hosts. Zhang *et al.* [28] identified the important *k*-mer features of viruses infecting the same host genera, and built a classifier to predict whether or not a new virus belongs to the same group of viruses. One drawback of the network-based approach is that the performance can diminish if the query virus is highly divergent from the known viruses in the current network.

Though various methods have been proposed for predicting virus-host interactions, the highest accuracy is only 50% at the genus level using a single type of information. With the increasing number of viruses being discovered, there is a demand for a tool that is able to accurately and rapidly predict the hosts of viruses, incorporating all types of virus-host and virus-virus features. In this paper, we have developed a network-based integrated framework for predicting virus-host interactions based on multiple types of information: virus-virus similarity, virus-host alignment-free similarity, virus-host shared CRISPR spacers and virus-host alignment-based matches. To the best of our knowledge, this is the first time that multiple types of features are effectively integrated into a stochastic network to complement each other and enhance the prediction accuracy of virus-host interactions. This integrated method markedly improved the accuracies in predicting virus-host interactions for complete viral genomes from 50% to 61% at the genus level, and yielded 85% accuracy at the phylum level, the highest among all the existing methods. The prediction framework also had decent accuracy for shorter viral contigs even as short as 10 kb. We have used our method to infer the host of 250 strains of crAss-like viruses, 1,811 marine viral genomes, and *>* 60,000 viral contigs from various environments. We have provided a user-friendly program, VirHostMatcher-Net, that uses this framework to predict virus-host interactions. Finally, VirHostMatcher-Net provides a flexible and expandable network-based framework for on-going refinement of virus-host prediction methods.

## Results

### A novel network-based integrated framework for predicting virus-host interaction

We collected from NCBI the genomes of a set of known virus-host interaction pairs, *S*_+_, and generated a set of random virus-host pairs that most likely do not interact, *S*_−_, as the training data for this study. Our objective was to develop a machine learning approach to predict the probability of interaction between a query virus-host pair (*v, b*), denoted as *P* (*I*(*v, b*) = 1), where *I*(*v, b*) denotes the interaction status of a virus *v* and a host *b* with value 1 indicating interaction and 0 indicating no interaction. In order to achieve the best performance, we comprehensively considered various factors that contribute to the interaction of a virus-host pair (*v, b*). First, if a virus is genetically close to viruses infecting a particular host, this virus is highly likely to infect the same host [28, 27]. On the other hand, if a virus infects a host, the virus should be genetically distant from the viruses that do not infect the host. Secondly, the similarity among hosts indicates the possibility of infection by the same virus [40, 41]. If a potential host belongs to the same taxon as the known host of the virus, then that host is likely to be infected by the virus. Third, the similarity between virus-host pairs in terms of genomic signatures reflects the likelihood of interaction [23]. If a virus genome is similar to a host genome in terms of the alignment-free *k*-mer usage pattern, the pair is predicted to have a high probability of interacting. Finally, the existence of virus-host shared CRISPR spacers and the alignment-based matches (i.e. BLASTn) are strong evidence of interaction.

All together, virus-virus similarity, host-host similarity, and virus-host similarity can be integrated to form a two-layer network connecting viruses and hosts. Thus, we constructed a virus-host pairs (VHP) network where nodes are VHPs and edge weights are the pairwise similarities between VHPs. We developed an integrated network-based Markov random field (MRF) approach that systematically and comprehensively integrates various types of features to predict interacting virus-host pairs. The probability of a given VHP to be interactive is based on the characteristics of this VHP itself, and the connectivity between this VHP and its neighbor VHPs in the network. Intuitively, the characteristics of a VHP itself include alignment-free score, the fraction of alignment-based matches, and the existence of shared CRISPR spacers. The connectivity between this VHP and other VHPs is defined based on the genome similarity between the virus and other viruses infecting the same host. The outline of the framework is demonstrated in Fig. 1. The details of the models for this framework can be found in the Methods section.

**Figure 1.**
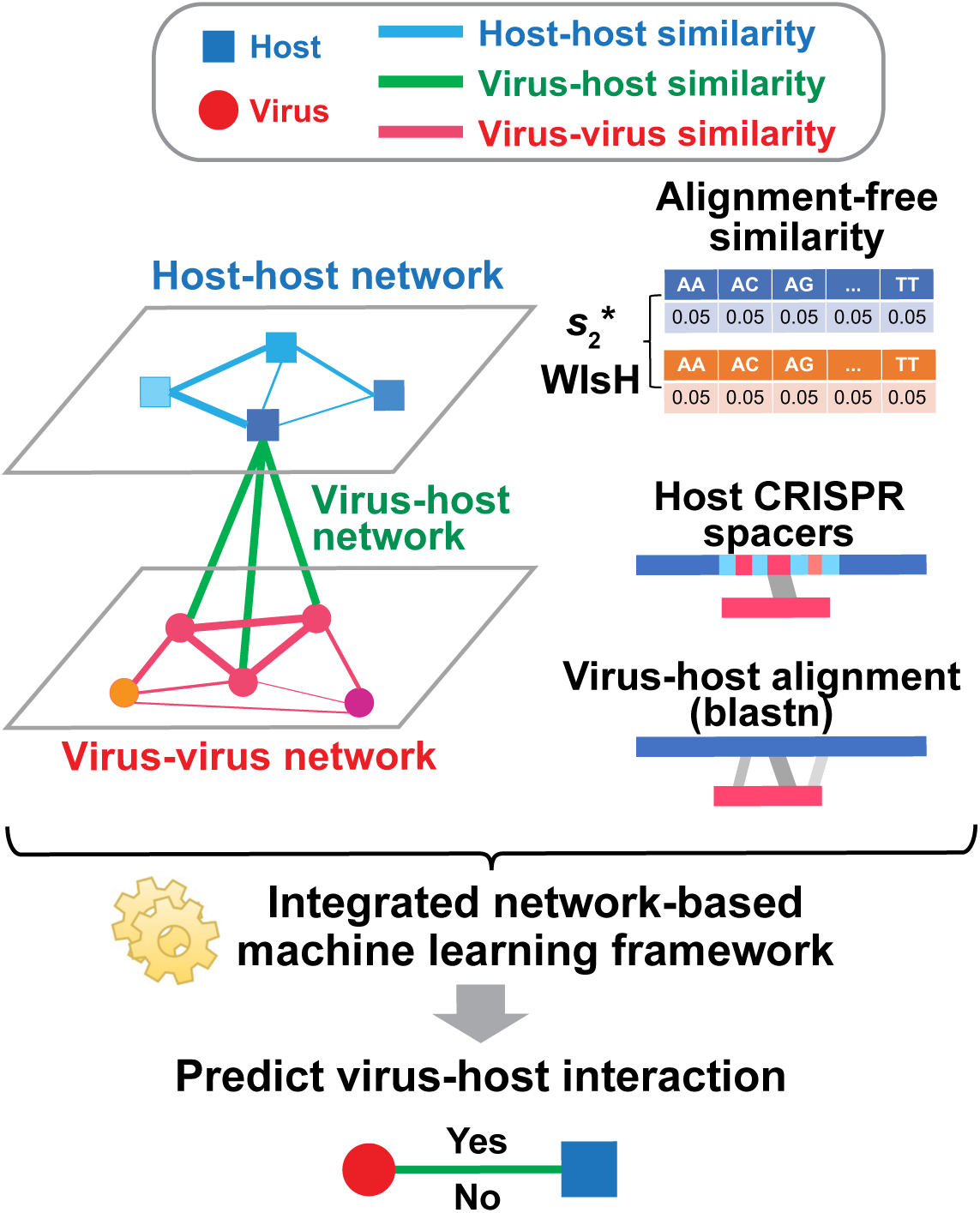
Overview of the network prediction framework. A novel two-layer network is constructed for representing virus-virus, host-host, and virus-host similarities. Viruses (red circles) are connected based on sequence similarity (red edges). Similarly, hosts (blue squares) are connected based on sequence similarity (blue edges). The thickness of the edges indicate the degree of similarity. The interaction between a virus and host pair (green edges) can be predicted using multiple types of features: 1) the similarity between the virus and other viruses infecting the host; 2) the similarity between the host and other hosts infected by the virus; 3) the alignment-free sequence similarity between the virus and the host based on *k*-mer frequencies; 4) the existence of shared CRISPR spacers between the virus and the host; 5) alignment-based sequence matches between the virus and the host. Finally, a network-based machine learning model is used to integrate all different types of features and to predict the likelihood of the interaction of a virus-host pair.

### Feature scores are significantly different between positive and negative virus-host pairs

We incorporated multiple types of features that contribute to the prediction of virus-host interactions. To assess the discriminatory power of each feature, we compared the distributions of the feature values between the virus-host interacting pairs and the non-interacting pairs. A set of 352 known virus-host interacting pairs was used as the positive set, and a set of the same number of randomly selected virus-host pairs was used as the negative set. See the Methods section for details of the data collection and the simulation of negative pairs. We used a one sided t-statistic to test if the feature values in the positive set are significantly higher or lower than the ones in the negative set.

First, the alignment-free similarity score 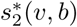 was used to measure the similarity between virus and host pairs, where 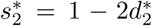 and the *k*-mer based dissimilarity score 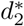 is defined in our previous work [23]. The measure 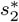 has an advantage over other classical similarity measures because of its precise correction of background noise, and has shown superior accuracy for predicting virus-host interactions [23]. See the Methods section for the definition of 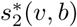. The 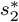 score had significantly higher values (*p*-value*<*2.2e-16, one sided t-test) for positive virus-host pairs than the negative pairs (Fig. 2a). The mean 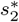 similarity score between positive pairs was 0.50 while the mean 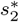 similarity between negative pairs was 0.27.

**Figure 2.**
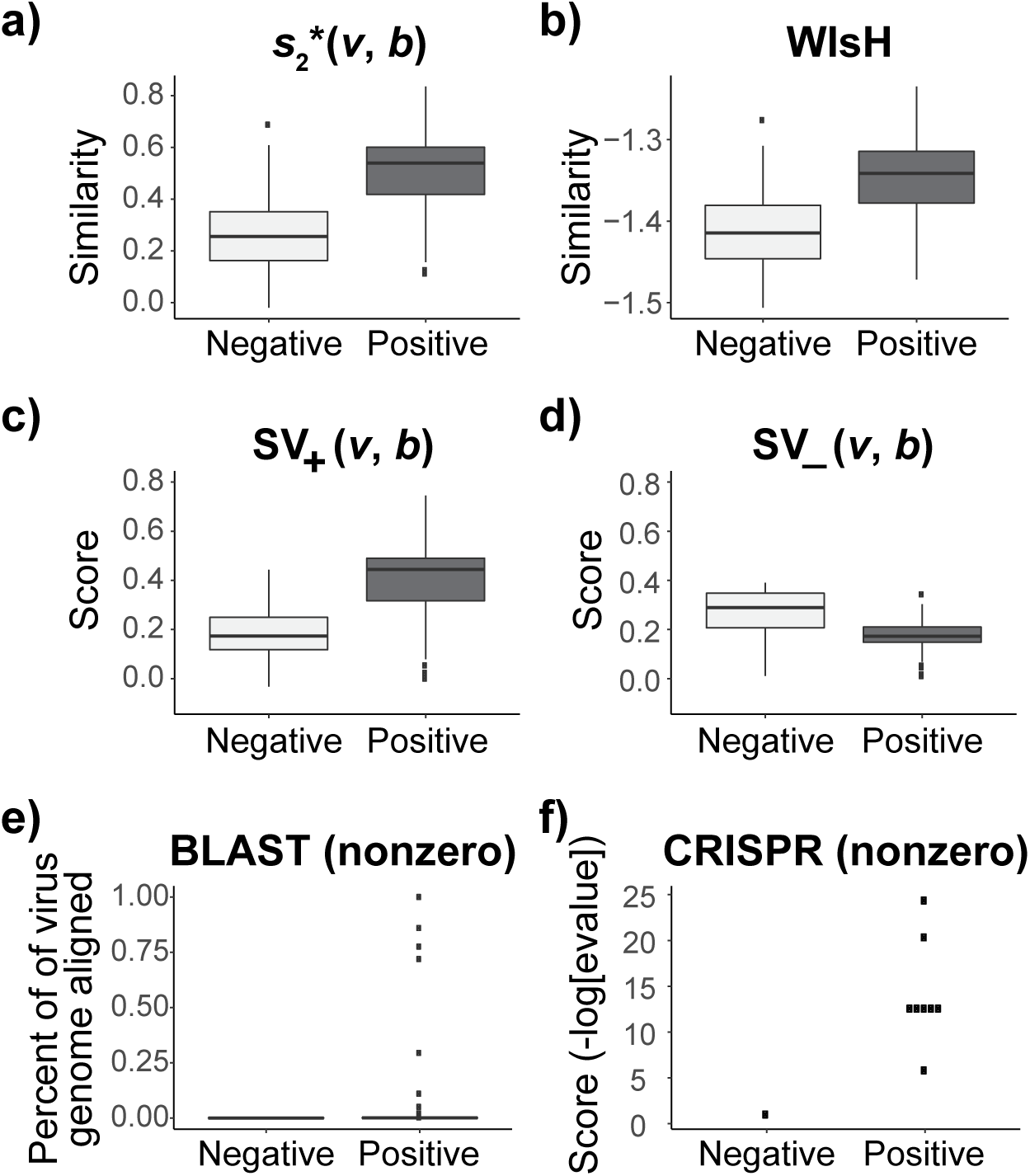
Distributions of the different feature values among 352 interacting and non-interacting virus-host pairs. The positive set consists of 352 known infecting virus-host pairs (positive set) and the same number of randomly selected virus and host pairs were used as the non-interacting, negative set. **a)** Boxplots of similarity defined by 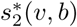. **b)** Boxplots of the log-likelihood scores given by WIsH; **c)** Boxplots of *SV*+(v, b) scores; **d)** Boxplots of the *SV*−(v, b) scores; **e)** Boxplots of BLAST scores. There are 108 nonzero values in the positive set and 79% of them are less than 0.01. **f)** Dotplots of the CRISPR scores. Only 8 virus-host pairs in the postive set and 1 pair in the negative set share CRISPR spacers. For boxplots in a-e, the horizontal bar displays the median; boxes display the first and third quartiles; whiskers depict minimum and maximum values; and points depict outliers beyond the whiskers.

The WIsH score, proposed by Galiez *et al.* [24], is another alignment-free similarity measure for a virus-host pair. It uses a log-likelihood score of a Markov chain model to measure similarity between viruses and hosts. We computed the WIsH scores for both positive and negative virus-host pairs, and found that the WIsH scores for positive virus-host interacting pairs is significantly higher than that for the negative virus-host pairs (*p*-value = 1e-10) (Fig. 2b). In fact, we observed that the WIsH and 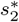 scores were highly correlated (Pearson correlation coefficient *ρ* = 0.83, *p*-value *<*2.2e-16). We predicted a virus-host pair as interacting if one of the similarity measures, 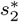 or WIsH, was above a threshold and, by changing the thresh-old, the corresponding receiver operating characteristic curve (ROC) was plotted. The area under the receiver operating characteristic curve (AUROC), which measures the discriminative ability between positive and negative pairs, was 0.90 for 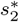 and 0.85 for WIsh (Additional File 1). Though the distinguishing power using WIsH was lower than that of 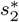 using complete genomes, WIsH was previously shown to be more effective than 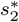 when predicting hosts of partial viral genomes [24]. Therefore, we decided to use 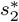 to measure virus-host alignment-free similarity when the length of viral sequence is close to the size of a complete genome, and to use WIsH to measure the virus-host similarity for short contigs.

Second, for a given virus-host pair (*v, b*), we defined the similarity between a virus *v* and other viruses infecting the host *b*, denoted as *SV*_+_(*v, b*), and likewise, the similarity between virus *v* and other viruses not infecting the host *b*, denoted as *SV*_−_(*v, b*). See Methods for the details of their definitions. We hypothesized that, for a true interacting virus-host pair (*v, b*), other viruses that infect the same host *b* should exhibit high similarity to the virus *v*, resulting in a high *SV*_+_(*v, b*). At the same time, other viruses not infecting the host *b* should have low similarity to the virus *v*, resulting a low *SV*_−_(*v, b*). For a non-interacting virus-host pair, the above trend of *SV*_+_(*v, b*) and *SV*_−_(*v, b*) should be opposite. Consistent with our hypothesis, *SV*_+_(*v, b*) scores were significantly higher for positive virus-host pairs than negative pairs, and vice versa for *SV*_−_(*v, b*) scores (both *p*-values *<*2.2e-16, Fig. 2c-d).

Third, we included information from CRISPR matches and alignment-based genome similarity between viruses and hosts. The CRISPR score was defined as the highest alignment score between the predicted CRISPR spacers in a host and a viral genome, and the alignment-based matching score was defined as the fraction of virus genome that significantly matches the host genome using blastn (*>* 90% identity, see Methods). Thus, for simplicity, we refer the alignment-based matching score to as the BLAST score. Both CRISPR and BLAST scores were significantly higher for the true interacting virus-host pairs than the non-interacting pairs with *p*-values of 0.004 and 0.0003 for one sided t-tests, respectively. Fig. 2e-f shows the limited frequency of CRISPR and BLAST matches between viruses and hosts. For the alignment-based BLAST score, only 108 (31%) out of 352 positive virus-host pairs had nonzero values, and 79% of the scores were less than 0.01. The CRISPR scores were even more sparse, with only eight nonzero values (2.3%) out of 352 in the positive set, which can explain why the *p*-value for comparing CRISPR scores was not as low as the other measures.

### Integrated approach markedly increases host prediction accuracy

We integrated the multiple types of features proposed previously to predict virus-host interactions using a general framework of MRF, where the nodes were virus-host pairs (VHP) and edges were the similarity between the VHPs. We investigated the prediction accuracies of the newly developed integrated models in equations (6) and (7) (see Methods), and compared the accuracies with those using the individual features. The model in Eq. (6) incorporates the network features including virus-virus similarity *SV*_+_ and *SV*_−_, the virus-host similarity 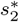, and the CRISPR score. The model in Eq. (7) combines features in Eq. (6) plus the BLAST scores. For each of the integrated models, we learned the parameters using the 352 positive and the same number of negative virus-host pairs, and then tested the trained model on the remaining 1,075 viruses for which their true hosts are known against 31,986 candidate hosts. The estimated coefficients and the corresponding *p*-values of the features are shown in Table 1. All the coefficients had the expected signs that were consistent with the observations in Fig. 2, and the statistical significance *p*-values for the coefficients were all *<*0.05. Note that although the *p*-value for *S*_*CRISPR*_ was 0.035, it was not as low as the *p*-values for the other features due to its limited availability.

**Table 1.**
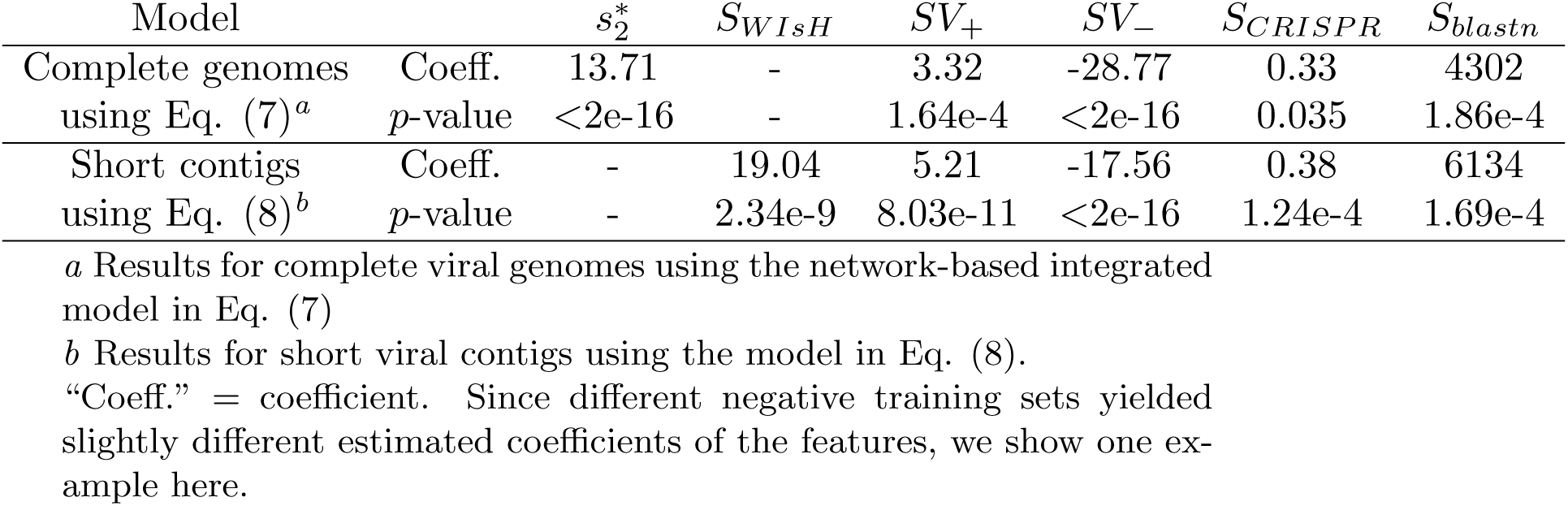
The estimated coefficients and corresponding *p*-values for host prediction features.

We assessed the prediction accuracies of the trained models using an independent set of 1,075 viruses at different taxonomic levels, including genus, family, order, class, and phylum. For each virus, we computed the prediction scores between this virus and all candidate hosts (*n* = 31, 986) using the trained models, and predicted the host as the one having the highest prediction score. The prediction accuracy was calculated as the percentage of viruses whose predicted hosts had the same taxonomy as their respective known hosts. Host prediction accuracies were markedly higher for the integrated approach using network features and CRISPR scores than using 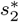 or CRISPR scores alone (Fig. 3a). For example, at the genus level, prediction accuracy was 34% and 36% when using 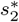 and CRISPR, respectively. Combining network similarity features together with CRISPR score (Eq.(6)) increased prediction accuracy to 54%, or a 1.5-fold increase.

**Figure 3.**
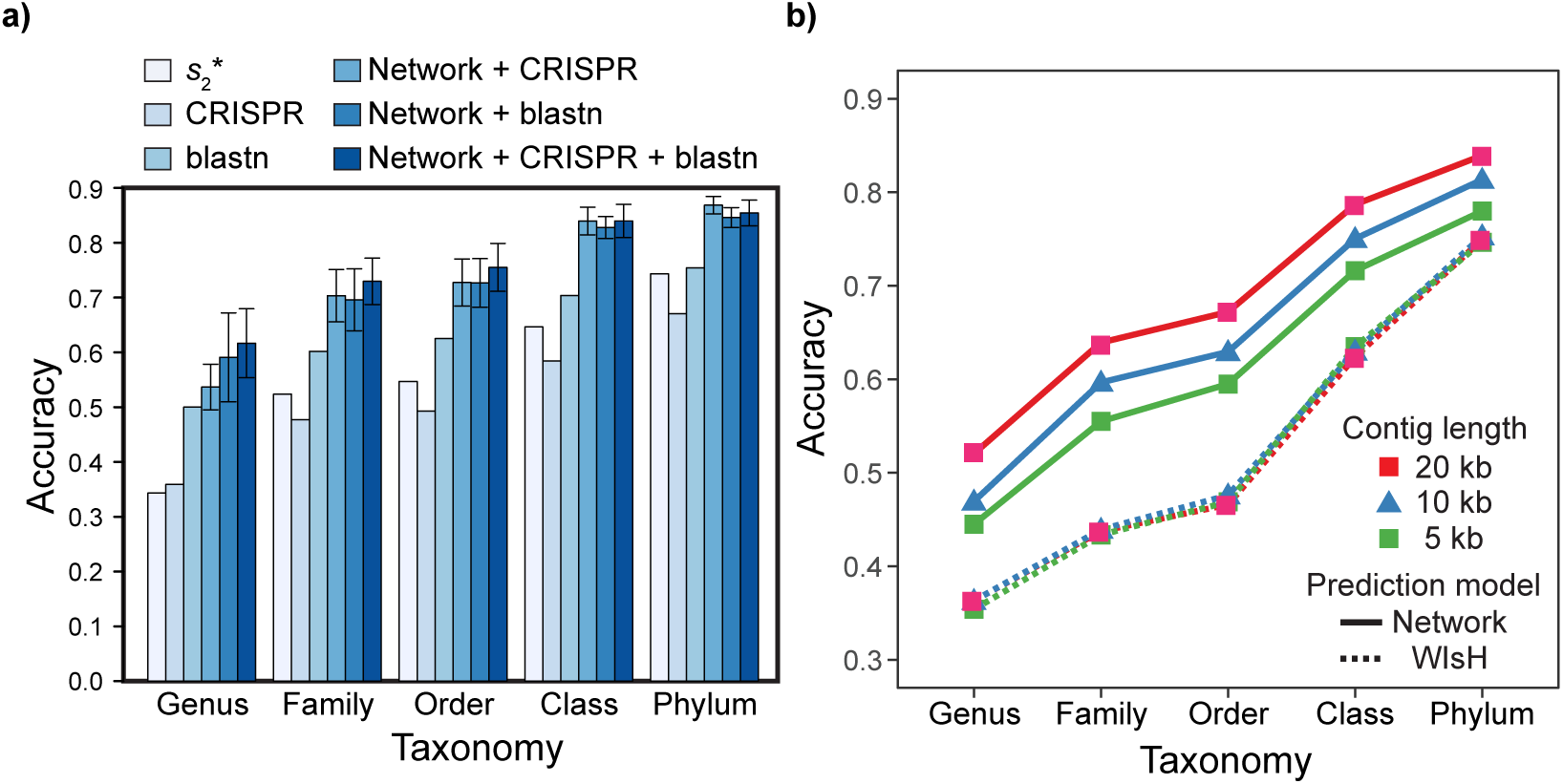
Prediction accuracies of the different approaches for 1,075 viruses (a) and simulated viral contigs (b). **a)** Prediction accuracies for 1,075 viral genomes whose true hosts are known against 31,986 candidate hosts, binned by taxonomic level. The first three bars show results using individual features of 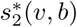, CRISPR score, or alignment-based similarity score (blastn), respectively. The remaining bars show results with integrated network models, trained using 352 positive and the same number of negative virus-host pairs as in Fig. 2. In order these are using Eq. (6) which incorporates the network-based features *SV*_+_(v,b) and *SV*_−_(v,b), alignment-free virus-host similarity 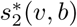, and the CRISPR match scores (“Network+CRISPR”); Eq. except that blastn scores are used in place of CRISPR scores (“Network+blastn”); and Eq. (7) that uses the features in Eq. (6) plus blastn scores (“Network+CRISPR+blastn”). Error bars for the network-based results depict 95% confidence intervals using 100 replicates of negative training sets (random virus-host pairs). **b)** Prediction accuracies for contigs subsampled at various lengths from the 1,075 virus genomes in **a)**. Mean accuracies are shown at different taxonomic levels using WIsH scores only (dashed lines) or the integrated model in Eq. (8) (solid line) that uses WIsH scores in place of of 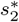 scores. Standard errors of prediction accuracies were determined for 100 replicates of subsampled contigs and were smaller than the plotted symbols, but for example at the genus level errors were 0.001, 0.001 and 0.002 for WIsH and 0.002, 0.002, 0.014 for the integrated model for contig lengths of 20 kb, 10 kb, and 5 kb, respectively.

Alignment-based BLAST scores alone had relatively high prediction accuracy. Incorporating BLAST scores into the network model by Eq. (7) increased genus-level host prediction accuracy to 62%, an additional 12% improvement. For the higher levels of taxonomy like family, order, class and phylum, the network-based integrated framework also achieved large improvements over the prediction accuracy of individual features, yielding 73%, 76%, 84%, and 85% prediction accuracy, respectively. Though both BLAST and CRISPR features helped improve the prediction accuracy, we noticed that the improvement made by adding the CRISPR score to the network model was higher, except at the genus level, than that of adding the BLAST score. This is interesting despite the apparent low occurrences of CRISPR sequences in our testing data set (Fig. 2). The model using all five features in Eq. (7) had the highest accuracy and was used in the subsequent host prediction applications. These results also demonstrate the great potential of our integrated framework: when other meaningful features contributing to virus-host interaction become available, the model is flexible such that it can be further extended to provide more accurate predictions.

### Integrated approach improves host prediction accuracy of short viral sequences

Viral contigs assembled from metagenomic data often represent partial viral genomes. We tested an integrated model in Eq. (8) that uses WIsH scores instead of 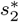 for measuring the alignment-free similarity between viruses and hosts. We evaluated the accuracy of the model for predicting the hosts of viral contigs at various lengths, and investigated the effect of viral sequence length on the prediction accuracy. To evaluate the performance of host prediction for short viral contigs, we randomly sub-sampled fragments of different lengths (5 kb, 10 kb, and 20 kb) from each of the 1,075 viral genomes. For a given viral genome and a fixed contig length, we randomly chose a segment of fixed length uniformly from the genome. If the fixed length was longer than the size of the complete genome, we took the entire genome. This procedure was repeated 30 times for each contig length. We then computed all the features of the contigs using the same procedure as for the complete viral genome analyses, with the only difference being that 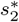 similarity was replaced with the WIsH score [24]. The model was trained with the same set of 352 virus-host positive pairs and the same number of negative pairs using the same scheme as before by replacing 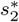 with the WIsH score. Similar to the model for predicting complete viral genomes, the estimated coefficients are shown in Table 1. With the trained model, we predicted the hosts for all sub-sampled contigs.

The average prediction accuracies for each contig length are shown in Fig. 3b. Our model (solid lines) achieved a large improvement compared to the results of WIsH alone (dashed lines). For example, when the contig length was 20 kb, the prediction accuracy using our model was about 16-21% higher than that of WIsH at the genus, family and order levels. As expected, the prediction accuracy of our model (solid lines) decreased with decreasing contig length. For instance, at the genus level, the accuracy decreased from 61% for the complete genome, to 52% for 20 kb, 47% for 10 kb, and 44% for 5 kb contigs. The standard error of the accuracy also increased as the contig length decreased. Note that the prediction accuracy of WIsH was not strongly affected by the contig length. Given the results, we provide our framework with two models for host prediction: one for complete or nearly complete viral genomes using the model in Eq. (7), and one for short viral contigs using the model in Eq. (8).

### Thresholding on the prediction score further improves accuracy

In many situations, investigators are interested in making sure the predicted hosts are as accurate as possible, i.e. the predictions have high precision or low false discovery rate. Therefore, we investigated how the accuracy changes by thresholding on the predicted probability of interaction *P* (*I*(*v, b*) = 1). In the above analysis, we predicted the host of every virus as the one with the highest score. However, sometimes the highest score was relatively low. For example, as shown in Fig. 4, the highest prediction score among the 31,986 hosts for some viruses in the complete genome test set was as low as 0.4. Low scores may occur, for example, when the true host is not in the database of potential hosts. In order to improve the prediction accuracy, we can set a threshold such that host predictions are only made if the score is above that threshold. For instance, when a threshold was set at 0.78, there was an improvement of prediction accuracy at all taxonomic levels. Specifically at the genus level, accuracy was improved by 8%, from 62% to 70%; at the phylum level, accuracy was improved by 5% from 85% to 90%. As a trade-off, the proportion of viruses whose hosts can be predicted among all the viruses (the recall rate) decreased. However, this tradeoff turned out to have minimal impact on the prediction result. In fact, when the threshold was set at 0.78, the recall rate only went down to 85%. The model only missed 3% of the viruses for which the model should have made correct predictions at the genus level. In practice, this small sacrifice can provide more confidence in predictions.

**Figure 4.**
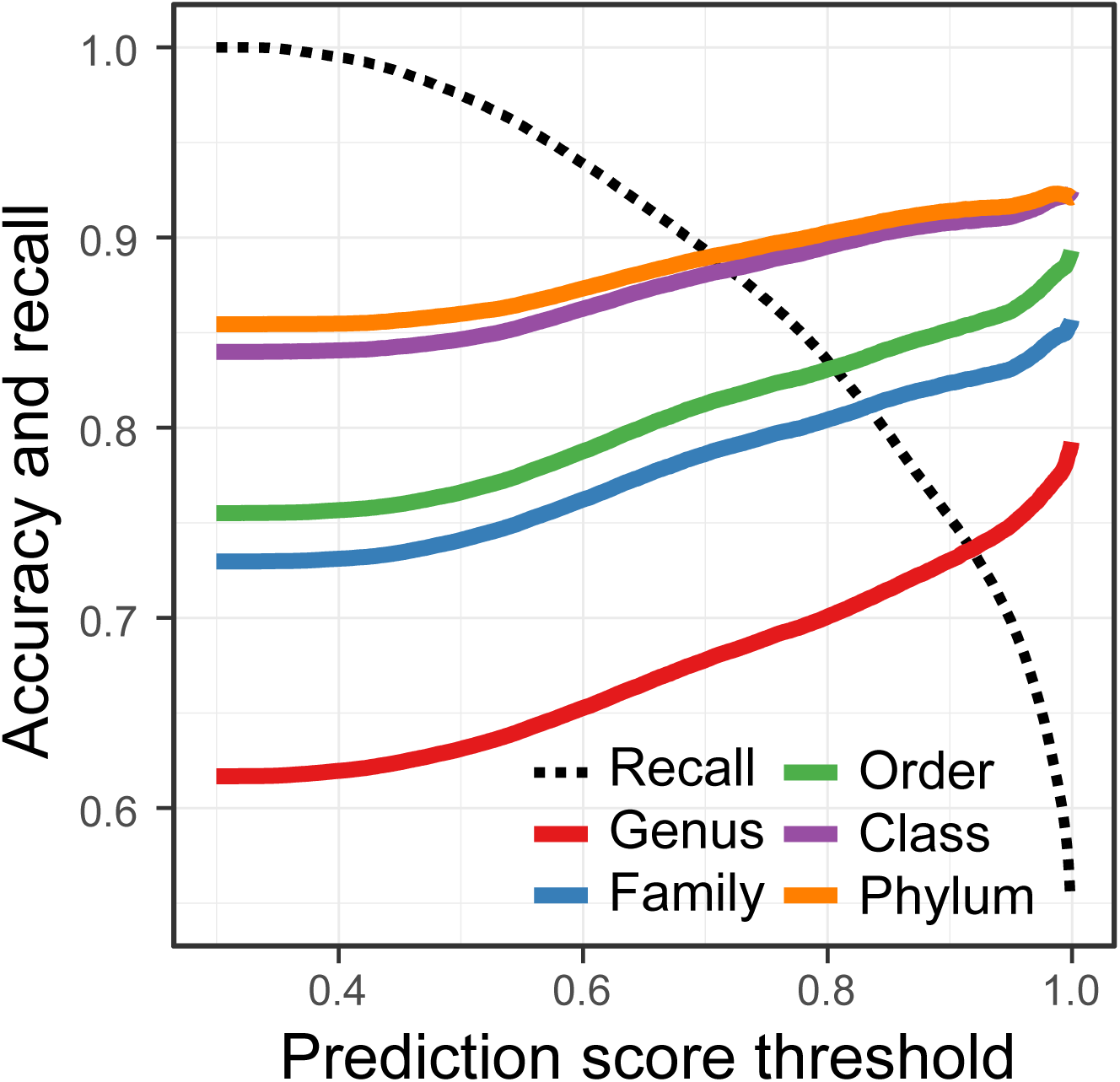
Improvement in host prediction by thresholding on the prediction score. By applying a given threshold, predictions were made only when the prediction score is above the threshold. Predictions were made using the whole genomes of 1,075 viruses whose true hosts are known among 31,986 hosts as in Fig. 3. The proportion of viruses that can be predicted (recall rate) decreases as the prediction accuracy at all levels increases.

### Prediction accuracy varies for different virus families

Viruses from three major families, *Siphoviridae, Myoviridae*, and *Podoviridae*, are highly represented in our evaluation data set. Previous host predictions with 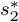 showed notable differences in prediction accuracy among these families [23]. Therefore, we examined prediction accuracies using our updated models (Fig. 5). We found that the *Siphoviridae* family of viruses in our data set had generally higher prediction accuracy than other families of viruses, achieving 73% accuracy compared with the average accuracy of 62% for all types of viruses, consistent with previous results using the 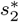 scores alone [23]. The prediction accuracies for the different virus families with various thresholds on the prediction score are shown in Additional File 2. We also noticed that the top prediction scores for the *Siphoviridae* family of viruses are significantly higher than that for the other two families (Kolmogorov-Smirnov test, *p*-value*<*1e-15). The above observations may be explained by the fact that siphoviruses typically have relatively narrow host ranges and podoviruses and myoviruses often have broader host ranges [42, 43, 44], though recent studies suggest that current isolation techniques may result in the under-representation of broad host range viruses and that the true host range of viruses is hard to define [40, 45].

**Figure 5.**
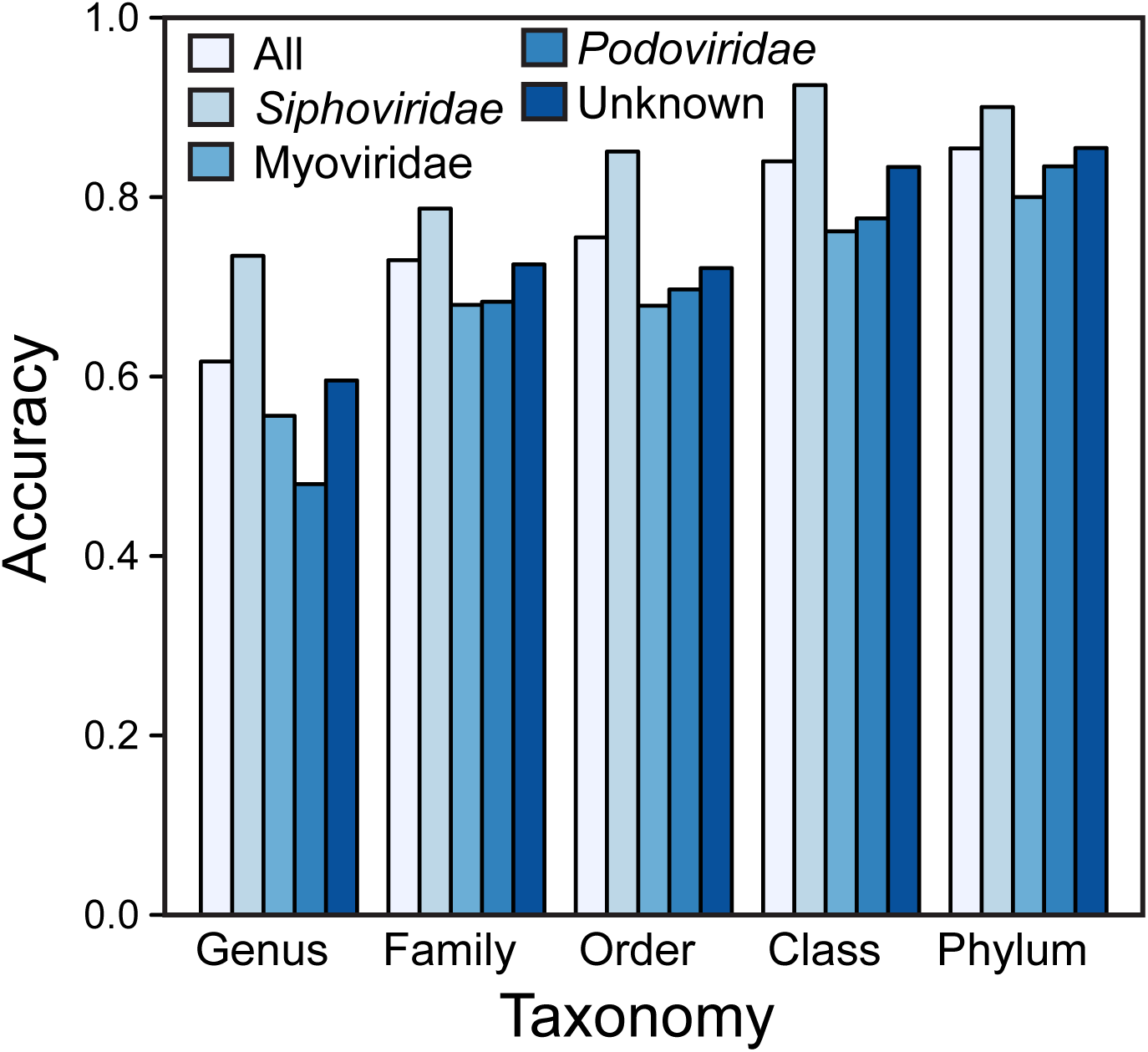
Differences in prediction accuracy across viral families. Prediction accuracies for different virus families within the order *Caudovirales*: *Siphoviridae, Myoviridae*, and *Podoviridae*. For comparison, accuracies are shown for all viruses (“all”) and for viruses outside of the *Caudovirales* or for which their virus families were not listed in the Genbank files (“other”). Predictions were made using whole viral genomes with no thresholding.

### Different genera of crAss-like phages infect a variety of hosts

CrAssphage was first discovered through the cross-assembly of human fecal metagenomes and was originally published as an individual genome that is referred as prototypical crAssphage (p-crAssphage) [19]. Though crAssphage is ubiquitous in human gut samples and comprises up to 90% of the sequencing reads in some fecal viral metagenomes [19], little is known about the biological significance and the hosts of crAssphage, due to the difficulty of culturing crAssphage and the high divergence between crAssphage and known viruses. Different methods have been used to predict the hosts of crAssphage. Dutilh *et al.* [19] predicted its host as the phylum Bacteroidetes using the co-occurrence profile between crAssphage and 404 potential human gut bacteria hosts across 151 human gut metagenomes from the Human Microbiome Project (HMP). Ahlgren *et al.* [23] compared the alignment-free similarity between crAssphage and the potential hosts, and the genera, *Bacteroides, Coprobacillus* and *Fusobacterium*, were found to have significantly high similarity to crAssphage.

Prediction of the host of crAss-like phage ΦcrAss001. Recent studies have found that the family of crAssphages is actually highly diverse. Two studies have recently used shared gene content to classify new and previously discovered crAss-like phage genomes from human gut viromes into four candidate subfamilies and 10 putative genera [46, 47]. Shkoporov *et al.* [48] isolated a particular strain of crAssphage, ΦcrAss001, by enriching viral fraction gut samples on a collection of 54 bacteria strains from the human gut. They subsequently showed that ΦcrAss001 specifically infects only one of 14 strains of *Bacteroides* tested, *Bacteroides intestinalis* 919/174. We first predicted the host of ΦcrAss001 using 24 species of the bacteria used to enrich ΦcrAss001 (and whose genomes are available) that span four phyla and 14 genera (Additional File 3). A *Bacteroides intestinalis* strain had the highest prediction score of 0.929, congruent with the experimental results of Shkoporov *et al.* [48]. Alignment-based scores such as CRISPR and BLAST were all 0 and did not contribute to the prediction. The main contribution comes from the alignment-free similarity score 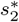 of 0.499. We then applied the integrated approach to predict the host of ΦcrAss001 using the large database of 31,986 host genomes and found that all of the top 25 predictions belong to the Bacteroidetes phylum, including 14 belonging to the genus *Prevotella*. ΦcrAss001 was classified as a genus VI crAssphage [48]. Guerin *et al.* [47] previously hypothesized that genus VI crAss-like phages infect *Prevotella* based on the observation that these two genera of virus and host were both enriched in malnourished and healthy Malawian infants. Our host prediction of ΦcrAss001 is therefore consistent with this hypothesis.

Prediction of the hosts of 249 crAss-like phages. We further predicted the hosts of 249 crAss-like phages in Guerin *et al.* [47] spanning all 10 of the putative genera of crAss-like phages against the full 31,986 host database (Additional File 4). All genus I crAss-like phages, including p-crAssphage, were predicted to infect *Akkermansia muciniphila*, in the phylum Verrucomicrobia, with the major contribution from BLAST scores. On average 5.1% of these phage genomes (∼5,000 bp) matched *Akkermansia muciniphila* with high similarity (97.2% identity on average). The aligned regions between p-crAssphage and *Akkermansia muciniphila* genomes ranged from 48 to 400 bp and were scattered across the p-crAssphage genome (Additional File 5). For viruses from the remaining crAssphage genera (*n* = 186), the majority of host predictions were to bacteria in the phyla Bacteroidetes (46%) or Firmicutes (35%), both major members of the human gut microbiome. In particular, *Prevotella* (phylum Bacteroidetes) was the most frequently predicted host genus (*n* = 34). crAss-like phages in genera VIII and IX were mostly found in Malawian gut samples, and as above these phage genera were hypothesized to also likely infect *Prevotella* [47]. Consistent with this suggestion, *>*80% of the predicted hosts were *Prevotella* for these two crAss-like phage genera using our method. A notable exception to these general prediction pattern of phyla Bacteroidetes or Firmicutes is genus II, for which most phages (9/14) had predicted hosts from the genus *Fusobacterium* in the phylum Fusobacteria. In particular, seven of these had their best match to *Fusobacterium perfoetens* ATCC 29250 isolated from pig feces [49]. The prediction scores for this crAssphage genus were generally lower than other genera, but were driven by high CRISPR scores. These results should be taken with some caution and investigated further.

### Host prediction for marine environmental viral genomes

Metagenomic sequencing has provided access to a broad range of viral genomes and has played an important role in studying uncharacterized marine viral genetic materials. Nishimura *et al.* [50] compiled a set of 1,811 marine environmental viral genomes (EVGs) including those newly assembled from the Tara Ocean [6] and Osaka Bay viromes and previously reported EVGs [51, 52, 53]. They predicted putative hosts of the EVGs based on the gene-based similarity between the EVGs and the cultured viral genomes with known hosts. In particular, they compiled another set of cultured viral genomes as a reference (RVGs) and created a proteomic tree for all EVGs and RVGs by the all-against-all distance matrix calculated from tBLASTx. They first assigned hosts by directly comparing the proteomic similarity between the EVGs and RVGs resulting in host assignment for 29 EVGs. They then constructed genus-level genomic operational taxonomic units (gOTUs) according to the proteomic tree. Based on the identification and phylogenetic analysis of various functional genes in EVGs and their closeness to related RVGs in the proteomic tree, they predicted the hosts of gOTUs at different host taxonomic levels (phylum to genus). In total they predicted the hosts for 564 EVGs.

We used our integrated model in Eq. (5) to predict the hosts for the 1,811 EVGs using a list of 3,528 marine hosts as host candidates that was provided in Ahlgren *et al.* [23]. We set a cutoff of 0.78 on the prediction score to ensure 90% prediction accuracy at the phylum level (Fig. 4). With this cutoff, our model was able to make host predictions for 1,239 EVGs, among which 347 EVGs also had phylum-level host predictions by Nishimura *et al.* (see Additional File 6 for the prediction results). Compared with the predictions of Nishimura *et al.*, our method had consistent predictions for 200 (58%) out of the EVGs at the phylum level and 168 (73%) out of the 229 EVGs at the class level (only 229 EVGs have predictions by our method and Nishimura *et al.*). In particular, our predictions were consistent with the previous predictions for the entire group of 23 cyanobacteria viruses. For a group of viruses that Nishimura *et al.* predicted to infect Proteobacteria, our predictions agree with theirs in 55 out of 75 cases at the phylum level. For another group of 201 viruses that were previously predicted as Flavobacteriaceae (within the phylum Bacteroidetes) phages, our predictions were consistent with theirs for 99 viruses at the family level. Note that the inconsistency between our predictions and Nishimura *et al.* may due to the different choices of features used for prediction. Predictions in Nishimura *et al.* are based on the similarity between virus genomes, while our method uses not only the similarity between viruses, but also the BLAST and CRISPR scores between virus and host genomes, which are direct evidence for interactions. In addition, our method was able to predict more hosts at lower taxonomic levels compared with the previous method. We had 322 of the 347 EVGs predicted at the order or lower host taxonomic level, a 13% increase in the number of EVGs that the previous method was able to predict.

For the 892 viruses whose hosts were not predicted previously and only predicted by our method, their predicted hosts include 13 phyla and 84 genera. In particular, we discovered 126 viruses infecting 59 novel host genera and 61 viruses infecting 6 novel host phyla that are absent from the data set of 1,427 isolate virus genomes (Additional File 6).

### Host prediction for metagenomic viral contigs from various habitats

Paez-Espino *et al.* [37] analyzed over three thousand geographically diverse metagenomic samples and identified 125,842 putative metagenomic viral contigs of median length 11 kb, revealing the extended viral genetic diversity in various environments [37]. In the original prediction, the metagenomic viral contigs and other 2,536 isolated contigs were first clustered into viral groups or singletons. They predicted the hosts of the viral contigs using a series of analyses including projecting the isolate viral-host information onto viral groups, matching viral contigs to a database of 3.5 million CRISPR spacers found in prokaryotic genomes, and identifying tRNA sequences in corresponding hosts. The analysis predicted hosts for 9,992 (7.7%) viral contigs. To evaluate our integrated approach for host prediction, we first used our method in Eq. (8) to predict the hosts of those putative metagenomic viral contigs. We then compared our predictions with those of Paez-Espino *et al.* by concentrating on 8,029 metagenomic contigs whose previously predicted host families were present in our host database and having a prediction score above 0.78. Our predictions were consistent with the vast majority of the original predictions, having 94% consistency at the phylum level (Table 2). Our predictions matched the previous predictions at an even higher rate (98% at the phylum level) for 53.1% of viruses whose hosts were previously inferred based on direct evidence of CRISPR spacer matches or tRNA matches to the hosts. For viruses whose hosts were inferred indirectly based on the hosts of other viruses in the same viral groups, our predicted hosts had 90% consistency with those based on the previous method at the phylum level. Thus, the inconsistent predictions mostly occurred for the viruses whose hosts were previously inferred based on viral group membership. For those viruses with inconsistent predictions, 66% of our predictions had significant network scores (*>*95% percentile), 44% had significant WIsH scores, 22% had significant CRISPR scores, and 22% had significant BLAST scores.

**Table 2.**
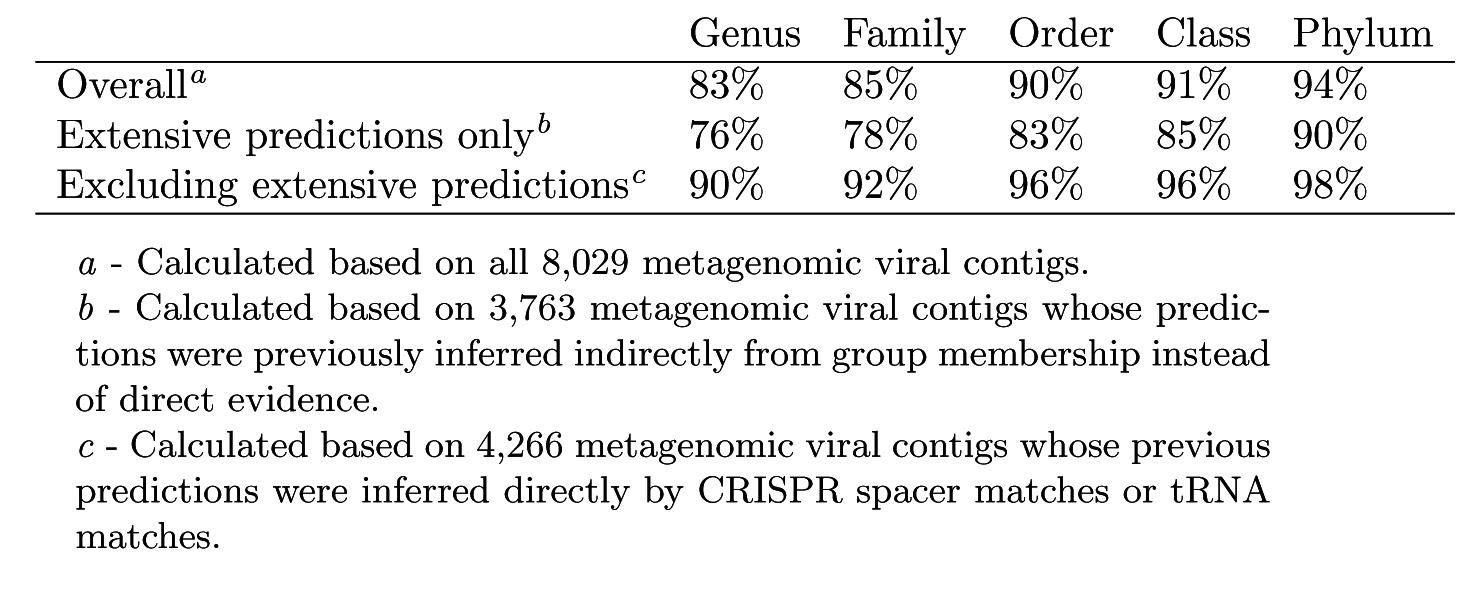
Proportions of congruent predictions for viral contigs between our method and those in Paez-Espino *et al.* [37]

We then predicted the hosts for the remaining available contigs that were not predicted in Paez-Espino *et al.* (*n*=101,343, note not all of the contigs from PaezEspino *et al.* are accessible at IMG/VR). Viruses were parsed by the type of sample from which they were obtained (human-associated, marine, and all other environments/sample types) and predictions were made against collections of host genomes corresponding to the sample type (HMP genomes, *n*=359; marine genomes, *n*=3,528; or all 31,986 host genomes, respectively). This resulted in 9,625, 31,041, and 20,643 viral contigs with prediction scores above 0.78 (Additional Files 7-9) or 61,309 viral contigs in sum. In combination with viral contigs with overlapping predictions by Paez-Espino *et al.*, we were able to make confident host predictions for 60% of all viral contigs, representing 6-fold more host predictions than previously by Paez-Espino *et al.*.

We then analyzed more specifically the predicted hosts for contigs with length ≥10 kb and for which ≥90% of their genes belong to known viral protein families (a criterion used in the original paper). There were 545 contigs from human associated samples that met the above criterion, and we restricted our host predictions to 359 HMP genomes. In total, 152 human-associated viral contigs were successfully predicted by our method with a score above 0.78 (Additional File 10). The predicted hosts of these 152 viral contigs belonged to 23 host genera. In particular, we discovered 59 viral contigs predicted to infect 15 host genera that have no known infecting viruses. To study the virus diversity within those hosts, we clustered the 68 viral contigs based on their percentage of shared genes using the UPGMA hierarchical clustering method. Viruses infecting the same host genus were mostly clustered into the same groups in most cases (Additional File 11). For example, viruses predicted to infect *Arcobacter, Lachnoclostridium, Blautia, Coprobacillus, Finegoldia* and *Ruminococcus* were grouped into clusters, respectively. On the other hand, viruses infecting *Fusobacterium* clustered into several groups, probably indicating a higher viral diversity in this host genus. It was also interesting to see that the viruses infecting the same host genus were from different samples in multiple studies (as assessed by contig IDs; WUGC, Baylor, LANL representing different studies), indicating those virus-host pairs were common across individuals. We also found that most of the newly identified pairs were habitat-specific. For example, the host genus *Arcobacter* and its 8 viruses were found in human tongue dorsum; *Lachnoclostridium* and its 5 viruses were in human stool; *Blautia* and its 3 viruses were in human stool; and *Veillonella* and its 5 viruses were in human tongue dorsum and throat. Though most virus-host pairs are present in specific habitats, we did notice exceptions such as the viruses infecting *Fusobacterium* in multiple human body sites like stool, tongue dorsum, supragingival, and throat. This may be consistent with the fact that *Fusobacterium* occurs in both the human gut [54] and oropharynx [55, 56]. Since *Fusobacterium* is a pathogen that causes oral and skin diseases, the newly discovered viral contigs infecting *Fusobacterium* may provide useful insights for phage therapy applications.

Similarly, we applied our method to a set of 558 marine viral contigs that were not predicted by Paez-Espino *et al.* using the same criteria as above. Prediction was restricted to the set of 3,528 marine hosts defined previously by Ahlgren *et al.*. Our model predicted hosts for 350 viral contigs using a score threshold of 0.78. The predicted hosts belonged to 13 host genera (Additional File 12). In particular, the newly identified virus-host pairs expanded the universe of known *Cellulophaga* viral diversity, a nascent marine heterotrophic model system. Previously, Holmfeldt et al. [57], by sequencing 31 viral isolates, demonstrated the existence of several viral genera associated with this marine group. Here we found an additional 160 viral contigs that putatively infect *Cellulophaga*. Using the same gene-based method for hierarchical clustering as in [57], the newly discovered 160 viruses clustered into multiple groups, including one having 49 contigs (group A) and one having 17 contigs (group B), which are separate from the group containing the 31 known isolates (Additional File 13). Overall, we identified at least 3 novel genera with each having more than 10 viral contigs, representing a sizable increase from the previously known diversity. Genera were defined, for consistency as in Holmfeldt et al., as pairs of genomes sharing more than 40% of their gene content. Those new virus groups were found in multiple locations such as Delaware Coast, Pacific Ocean and North Sea, indicating their ubiquity and potential impacts on communities of *Cellulophaga*, an important degrader of complex organic matter. In addition, our method predicted 140 viruses as cyanobacterial phages (cyanophages) infecting *Prochlorococcus*, a group of globally abundant marine cyanobacteria [58]. We independently confirmed that 71 of these are actually cyanophages based on significant nucleotide similarity to cyanophage isolate genomes (BLASTN, ≥65% identity over ≥40% of the contigs or ANI values ≥70%). The remaining 69 contigs thus represent potentially novel lineages that have no significant nucleotide similarity to known cyanophage isolates. This showcases both the diversity of virus-host interactions and the power of our method to capture groups with relatively few known representatives.

## Discussion

The interactions between virus and prokaryotic hosts play important roles in human health and ecosystems. Millions of new viruses have been identified using high-throughput metagenomic sequencing technologies, but little is known about their biological functions and the prokaryotic hosts with which they interact. We developed a network-based integrated framework for predicting the hosts of prokaryotic viruses. The new method provides a sizable improvement on prediction accuracy compared with previous methods by integrating multiple measures for informing host prediction. Based on the evaluation of the methods using a large benchmark data set containing 1,075 viruses and 31,986 hosts, the method achieves 62% and 85% prediction accuracy at the genus and the phylum levels, respectively, yielding 12% and 10% improvements at the genus and the phylum levels compared to the highest accuracy achieved by previous methods.

The novel two-layer network of virus-virus, host-host, and virus-host genomic similarity lays the foundation for this method. The employment of a two-layer network is inspired by underlying biological phenomena. First, it is observed that genetically similar viruses tend to infect closely related hosts [40, 41]. So the host of a new virus can be partly inferred based on the similarity to related viruses with known hosts. Similarly, the host of new viruses could potentially be inferred through similarity of hosts. Second, because viruses depend on the cellular machinery of their host to replicate, viruses often share highly similar patterns in codon usage or short nucleotide words with their hosts. The host of a new virus can be predicted using nucleotide word similarity between the virus and candidate hosts [11, 22, 23]. Thus, the two-layer network model is a natural formulation of the biological relationships described above. Despite the fact that the viruses in our current database only have one reported host for each virus such that host-host network connections cannot be incorporated into the prediction model, the novel two-layer network can be fully implemented in the future as multiple hosts of viruses are revealed.

Multiple types of features, including shared sequences between host CRISPR spacers and viral genomes and virus-host BLAST matches, combined with the network-based features, were incorporated to obtain an integrated framework for host prediction. The CRISPR and BLAST features are based on the biological process that some viruses and their hosts share a portion of their genomes due to CRISPR defence system, horizontal gene transfer, or prophage integration. Although these features have been investigated individually in previous studies [22, 23, 24, 35], this is the first time that multiple types of features have been integrated into a unified framework for virus-host prediction. Our results show that the integrated method combining all features achieves the highest prediction accuracy.

Our model also markedly improved the host prediction accuracy on shorter viral fragments at all taxonomic levels when compared to WIsH [24], a recently developed probabilistic method for predicting hosts of viral contigs. Our method was able to obtain 52%, 47% and 44% prediction accuracies at the genus level for 20 kb, 10 kb and 5 kb sequence lengths, respectively. The prediction accuracies for 20 kb, 10 kb, and 5 kb contigs were all above 78% at the phylum level. In practice, we recommend using a contig length of at least 10 kb for host prediction to ensure prediction accuracy greater than 80% at the phylum level.

Setting a minimum threshold for making predictions led to a notable improvement in accuracy and we show that this improvement comes at a minimal cost in recall. We also investigated the host prediction accuracy for different groups of viruses. Specifically, our observations indicate that viruses in the *Siphoviridae* family have higher prediction accuracy than the other *Caudovirales* families, consistent with the fact that siphoviruses tend to have a narrower range of target hosts [43, 44]. Likewise, restricting the possible hosts from all available prokaryotic genomes to a focused set of relevant microbes can help improve prediction accuracy, as was the case of predicting hosts of human associated viruses using the 359 HMP genomes and predicting marine viruses using 3,528 marine host genomes.

We utilized our model to predict the hosts of a family of crAss-like phages, a ubiquitous, abundant and highly diverse group of human gut viruses. A new strain of crAss-like phages, ΦcrAss001, was recently isolated and was found to infect *Bacteroides intestinalis* among a set of 54 strains belonging to 24 bacterial species [48]. Our computational prediction for the host of ΦcrAss001 against the 24 species for which genomes were available is consistent with the culture-based results. When we predicted its host against the 31,986 candidate genomes, the genus *Prevotella* within the Bacteroidetes phylum was the top predicted host. Although this genus is different from the experimentally determined host, the prediction of *Prevotella* is consistent with the hypothesis of Guerin *et al.* [47] that genus VI crAss-like phages, to which crAss001 belongs, infect *Prevotella*. We further predicted the hosts of 249 other crAss-like phages and the major host phyla include Bacteroidetes, Firmicutes, and Verrucomicrobia. Host prediction results also suggest that genera VII and IX crAss-like phages infect *Prevotella*, which again is consistent with patterns that these genera of viruses and *Prevotella* are together enriched in particular human populations. In particular, genus I crAss-like phages share roughly 5% of their genomes with the Firmicute *Akkermansia muciniphila*, strongly indicating that they interact with this host. Co-abundance correlation analysis of genus I crAss-like phage and host genera in HMP samples also supported *Akkermansia muciniphila* as a likely host, more strongly so than any other host genus, including *Bacteroides*. As additional evidence to support this discovery, we computed the Spearman’s rank correlation coefficient between the abundance profiles of p-crAssphage, a member of the genus I crAss-like phages, and *Akkermansia muciniphila* in 151 HMP samples that were used in a previous study [19]. We excluded samples that did not have any reads mapped to p-crAssphage by assuming *Akkermansia muciniphila* could be the host of other viruses. The coefficient between p-crAssphage and *Akkermansia muciniphila* was 0.27 (*p*-value = 0.008), higher than that for other human gut bacteria strains including *Bacteroides* strains. These combined results provide compelling evidence to further investigate *Akkermansia muciniphila* as a potential host of genus I crAss-like phages. As an increasing number of studies show the potential role that *Akkermansia muciniphila* plays in inflammation, obesity, type 2 diabetes, and the efficacy of immunotherapy [59, 60, 61, 62], further experimental validation for the interaction between genus I crAss-like phages and *Akkermansia muciniphila* is warranted. Overall, our predictions for crAss-like phages are consistent with previous experimental evidence [48] and previous hypotheses in Guerin *et al.* [47], including the suggestion that these viruses infect diverse hosts.

We also applied our method to predict hosts for viruses in two large-scale metagenomic data sets, one focusing on marine viral genomes such as those discovered in Tara Oceans, and the other including viral contigs in over three thousand geographically diverse metagenomic samples including marine and HMP samples. Our predictions had high consistency with previous predictions made using simple methods such as CRISPR or tRNA matches or gene-based similarity to known reference viruses. More importantly, our method greatly increased the number of viruses for which predictions could be made, six-fold more viruses than by Paez-Espino et al. These predictions were made using a minimum score threshold of 0.78, with a false discovery rate of *<*10% for nearly complete genomes and *<*20% for contigs of length *>* 10 kb at the phylum level. The newly predicted virus-host pairs revealed viruses for hosts without known infecting viruses, and also expanded the diversity of viruses for hosts with known isolate viruses, showcasing the usefulness of our method in expanding knowledge of hosts in both ways.

A major advantage of our network-based integrated framework is that it can be easily extended to incorporate more meaningful features that can better inform virus-host interactions in the future. Virus-host co-abundance profiles have been shown to provide some evidence of virus-host interactions [63, 64], but Edwards *et al.* [22] suggested that its performance on host prediction was relatively poor compared to other measures such as CRISPR and sequence homology. Coenen et al. [65] also showed that virus-host correlations are poor predictors of virus-host interactions. Our preliminary analysis of incorporating such co-abundance data as a feature likewise showed the model did not benefit from adding the co-abundance feature (see Additional File 14). In general, co-abundance can be a misleading feature because virus-host interactions may not always yield positive or negative correlations depending on the complexity of virus lifestyles (e.g. lytic vs. lysogenic) [66]. In fact, we noticed that the feature coefficient for co-abundance when incorporated into the model was not statistically significant, indicating that the co-abundance can not consistently be a useful predictor. Moreover, virus-host interactions are dynamic with delays and fluctuate over time, while metagenomic sampling only captures the community at a single time point. Also the interactions can be nonlinear because of the complicated many-to-many virus-host networks [65]. Likewise non-specific hosts and viruses can exhibit spurious correlations due to the computational bias in terms of the compositional data where the abundance vector is constrained to a constant sum. Similarly, hosts may be incorrectly predicted to infect certain viruses because their hosts coincidentally share similar niches and dynamics. Significant co-abundance between a virus and a host nonetheless is consistent with and can support in some cases the discovery of a true virus-host interaction, but co-abundance evidence alone should be taken with caution. Although we do not exclude the possibility that co-abundance could be useful under certain environments or for certain types of viruses, it is not likely that a simple co-abundance measure based on non-time series samples can well describe the virus-host dynamics in general. If other promising predictive virus-host features are discovered in the future, these can easily be incorporated into our framework.

Sequence-based and alignment-based measures such as CRISPR and BLAST scores generally have limited availability, but can provide solid evidence for virus-host interactions when such signals are present. On the other hand, alignment-free 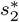 similarity can be computed for any virus-host pairs, but may not always perform as well as CRISPR and BLAST. We compared the prediction accuracies for 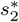 score and BLAST score when the hosts belonging to the true host genus of the viruses are removed from the candidates. The result showed that when the specific hosts were removed, the prediction accuracy for BLAST at the family level decreased markedly to 0.20, while the accuracy for 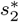 was 0.32 (Additional File 15). Therefore, alignment-based methods depend heavily on the existence of the true host in the database, and they can perform much worse than the alignment-free based methods for predicting hosts of new viruses when the true host genus is not in the host candidate set. These results highlight again how the integrated framework combining both alignment and alignment-free based features helps to complement the two types of methods and improve the overall prediction accuracy.

Although the new model makes sizable improvements over existing methods for both complete viral genomes and viral contigs at different taxonomic levels, the prediction accuracy at the genus level is still 62% for complete genomes and 50% for 10kb contigs. It is expected that with an increased data set of hosts and virus-host interactions for training our models, the prediction accuracy of our method will further increase. Our host data set will be gradually updated to include more newly discovered virus-host pairs for training and testing. However, we note that prediction accuracy at the phylum level is already very high (∼90%). Since there are many prokaryotic phyla (*>*75%) for which their viruses have yet to be identified, our tool is promising to greatly expand characterization of novel groups of viruses.

## Conclusions

In summary, our novel network-based integrated approach demonstrates how integration of multiple features informative of virus-host interactions significantly improves host prediction than any single feature. Application of our method to a few datasets of metagenomically assembled contigs demonstrate on the strong prediction ability of the model–yielding predictions largely congruent with previous methods but more importantly generating many more host predictions and identifying novel virus-host interactions than previous approaches. This approach will be valuable for identifying the putative hosts of newly discovered viral genomes particularly for the flood of new viral metagenomic data currently being generated. The flexible nature of our prediction framework also has the potential to be updated as new computational theories and biological understanding in virus-host interactions become available.

## Methods

### Data sets

All data generated or analyzed during this study are available from previously published studies or are included in this published article and its supplementary information files as detailed below. We used the same data as in [23] that includes 1,427 RefSeq viral genomes and 31,986 prokaryotic (bacterial and archaeal) genomes downloaded from NCBI on 5/8/2015. The viral data set includes all viruses with known hosts at the genus level at that time. The hosts of the viruses from which the viruses were originally isolated were collected based on the key words ‘isolate host=’ or ‘host=’ within each Genbank file. Furthermore, for a subset of 352 viral genomes, their hosts were reported at either strain, subspecies, or serovar, and only a single host genome was reported in the NCBI genome database for that particular strain, subspecies or serovar. We used the 352 viruses with known specific host genomes as the training set. The other viruses have host taxonomic information only down to the genus level. The accession numbers of all viruses and their hosts can be found in the supplementary material of Ahlgren *et al.* [23].

We used the developed computational method to predict the hosts of a family of 250 crAssphages [19], a newly discovered family of viruses from the human gut samples. These viruses are abundant and ubiquitous in human gut.

We also applied our method to a set of 1,811 marine virus genomes that were studied in Nishimura et al. [50]. The data set is available from [67]. In addition, we predicted the hosts of 69,338 viral contigs that were assembled previously from various environmental metagenomic samples [37]. Accession numbers of those viral contigs are available in Table S19 of Paez-Espino et al. [37]. The accession numbers for the novel virus-host pairs predicted exclusively by our method can be found in Additional Files 7-9.

### Outline of the model

We formulate the virus-host interactions using a Markov random field (MRF) model [29, 68]. Given a set of viruses *{v*_1_, *v*_2_, *…, v*_*n*_*}* and a set of hosts *{b*_1_, *b*_2_, *…, b*_*m*_*}*, we define the set of virus-host pairs (VHP) and their interaction statuses,

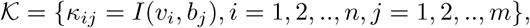

where *I*(*v, b*) = 1 if *v* infects *b* and *I*(*v, b*) = 0 otherwise. We construct a VHP network where nodes are VHPs and edge weights are the pairwise similarity between two VHPs.

The interaction statuses of all VHPs depend on two essential components: 1) the linkage between each VHP and all others and 2) the likelihood of the interaction status of each individual VHP. In the following sections, we first show how a MRF model can take the first component into consideration. Next we introduce the similarity measure 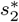 that describes the linkage between VHPs. Then we define all other features that can be used to estimate the second component. Finally, we derive two models for host prediction given virus genomes and contigs, respectively.

### A MRF approach for virus-host interactions

The likelihood of an assignment *𝒦* of the infection statuses for all the VHPs in the network is proportional to the likelihood of the assignments of the VHP nodes and the likelihood of the pairwise labels of VHPs given the network. Let *π* be the probability for a VHP to interact. Then for each pair (*v*_*i*_, *b*_*j*_), the likelihood of the interaction status, *P* (*𝒦*_*ij*_ = *κ*_*ij*_), can be expressed as 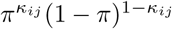 according to the Bernoulli model. By considering all VHPs and assuming their assignments are independent, the likelihood of an assignment of *𝒦* equals to the product of the likelihood for all the virus-host pairs, that is,

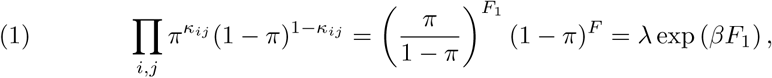

where *F*_1_ = Σ*κ* _*ij*_, *F* = ‖ 𝒦 ‖ is the size of, 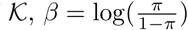, and λ = (1 − *π*)^*F*^.

Next consider the relationship between two VHPs in the network. The probability of two similar VHPs having the same 0-1 status is higher than the probability of having different 0-1 assignments. Let S_*ij,i*′*j*′_ be the similarity between two VHPs (*v*_*i*_, *b*_*j*_) and (*v*_*i*′_, *b*_*j*′_). Conditional on the similarity between two VHPs, we model the probability for them to be labelled as (1,1), (1,0) and (0,0) by 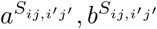 and 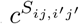, respectively, where *a, b*, and *c* are parameters. Mathematically, we can write the probability of (*v*_*i*_, *b*_*j*_) labeled as *κ*_*ij*_ and (*v*_*i*′_, *b*_*j*′_) labeled as *κ*_*i*′*j*′_ by

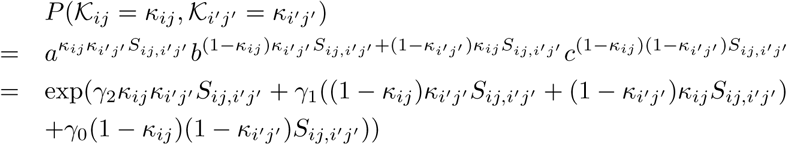

where *γ*_2_ = log(*a*), *γ*_1_ = log(*b*), and *γ*_0_ = log(*c*). We assume that the labeling of the VHP pairs are independent. Then we can multiply the above equation over all the VHP pairs to obtain

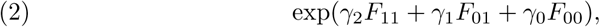

where *F*_*cc*′_ is defined as the sum of similarities among VHP pairs labeled as (*c, c*′), *c, c*′ = 0, 1, namely

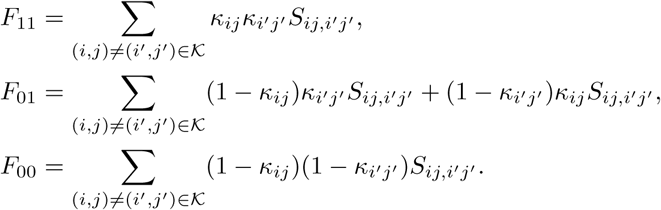

By multiplying equations 1 and 2 and then normalizing to a probability distribution, we model the probability of the assignment conditional on the similarity network as

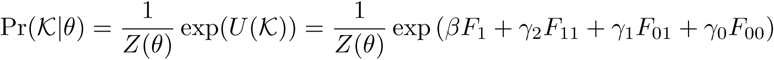

where *θ* = (*β*, *γ*_2_, *γ*_1_, *γ*_0_) are the parameters, and *Z*(*θ*) is the normalizing factor.

With this distribution function, for any *κ*_*ij*_ ∈ *𝒦*, we can calculate

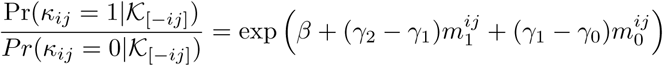

where

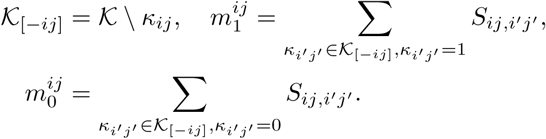

Then the log-odds of the probability Pr(*κ*_*ij*_ = 1| *𝒦*_[−*ij*]_, *θ*) is

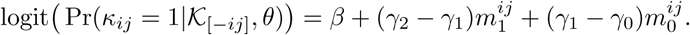

Denote *γ*_+_ = *γ*_2_ − *γ*_1_ and *γ*_−_ = *γ*_1_ − *γ*_0_. Then we have

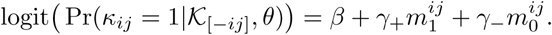

### The similarity between two VHPs and the generalized probability model for a VHP to interact

The MRF network model is constructed based on the similarity between pairs of VHPs *S*_*ij,i′j′*_. Various similarity measures between VHPs can be defined. In this study, we define the similarity between two VHPs as the similarity between the two viruses plus the similarity between the two hosts. To measure the similarity between two genomic sequences, we previously developed dissimilarity measures 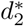 and 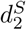 for alignment-free sequence comparison using *k*-mers as genomic signatures [69, 70, 71, 72], and showed that the dissimilarity measure 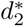 and 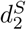 have high correlation with alignment-based distance measures [73]. Since viruses are highly diverse and alignments of highly divergent sequences are challenging, alignment-free measures are more suitable for sequence comparison than the alignment-based methods. Furthermore, Ahlgren *et al.* [23] showed that 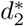 outperformed 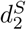 for the comparison of virus and bacterial sequences for the purpose of virus-host interaction prediction. Therefore, here we choose to use 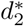 and transform it to 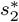 to measure the similarity between two sequences.

For each sequence, we represent it by the normalized *k*-mer frequency vector 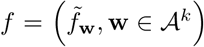, where *𝒜* is the set of alphabet {*A, C, G, T*}, *k* is the length of *k*-mer,

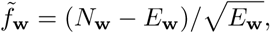

with *N*_**w**_ and *E*_**w**_ being the observed and expected numbers of occurrences of word **w** in the sequence. The expected count is calculated under a certain Markov chain model for the sequence. Since it was shown in [23] that *k* = 6 and second order Markov chain performed well in virus-host interaction prediction, we choose *k* = 6 and second order Markov chain in this study. The similarity between two sequences, 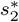, is defined as the un-centered correlation between their corresponding normalized frequency vectors. That is,

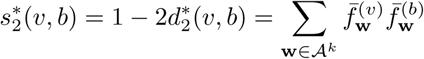

where 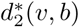 is the dissimilarity measure used in the previous studies, and 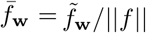 with ‖*f* ‖ being the Euclid norm of the feature vector 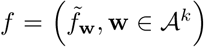 and the superscript indicates the virus The above formulation takes into accoun*v* or bacterial *b* sequence. Thus, we define the similarity

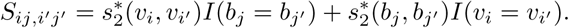

Plugging *S*_*ij,i*′ *j*′_ into the logit function, we have

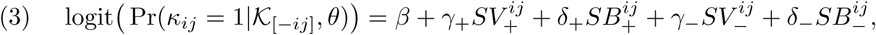

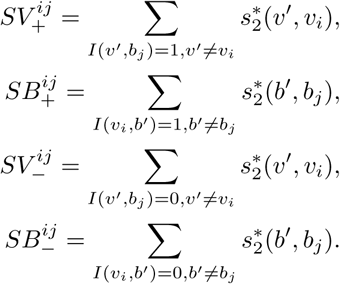

The above formulation takes into account both the similarity network between viruses, and the similarity network between hosts. In our data set, however, each virus has only one reported host. So when we train the model using the current data set, both 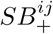 and 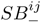 are set to zero. Then the model reduces to,

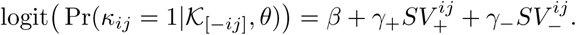

Though the terms 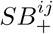 and 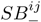 cannot be used given the current data set, as more virus-host pairs are collected in the training data, the host-host similarity network will contribute to the prediction model and the two-layer MRF network will be fully utilized based on Eq. (3).

#### Incorporating similarity between virus and host for interaction prediction

The assumption that any VHP has the same probability *π* for interaction is not realistic. Different pairs of virus and host have different features that affect the probability of interaction. For example, the probability can be associated with the similarity between the virus and the host. Thus, the probability *π* is modelled specifically to each individual pair (*v*_*i*_, *b*_*j*_),

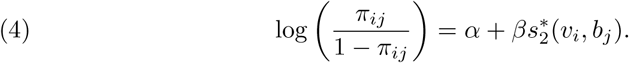

Then the logit model with the generalized probability can be written as,

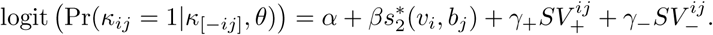

Therefore, the network-based MRF for predicting virus-host interaction is finally written as a logistic regression model where the predictors are the features of virus-virus similarity and virus-host similarity,

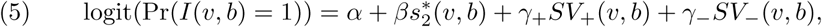

where *α* is a constant, (*β*, *γ*_+_, *γ*_−_) measure the contributions of the features 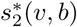, and *SV*_−_(*v, b*), respectively. We expect that *β* and *γ*_+_ to be positive and *γ*_−_ to be negative. However, we do not make these assumptions and let the data inform us the values of these parameters. To learn the parameters, we trained the model in a smaller training data set, and predicted virus-host interactions in the network of all viruses and hosts. Since the scales of *SV*_+_(*v, b*) and *SV*_−_(*v, b*) are proportional to the size of the data set, in practice we used the normalized variables, that is,

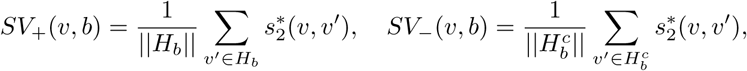

where 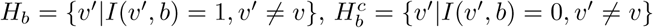, and ‖ · ‖ is the size of the set. When *H*_*b*_ or 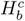 is an empty set, the value of *SV*_+_(*v, b*) or *SV*_−_(*v, b*) is set to zero.

To achieve the best performance, in addition to the similarity score 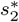, we integrate other types of features, including the CRISPR score and the alignment score between the virus *v* and host *b* into the framework.

### Sharing of CRISPR spacers between the virus and the host

The CRISPR systems play an important role as an adaptive and heritable immune system for prokaryotes. They help the host fight against the invasion of specific viruses by inserting small fragments of viral genomes (typically 21-72bp) as spacers into a CRISPR locus. The spacers are transcribed and are used as a guide by a Cas complex to target the degradation of the corresponding viral DNA [74].

Given a host genome, the CRISPR locus can be computationally located and thus the spacers can be extracted. In our study, we used the CRISPR Recognition Tool (CRT) [75] to find spacers. The spacers in a host genome (if available) were aligned to a viral genome by `blastn` [76] and alignment with E-value less than 1 were recorded. This threshold was chosen the same as the one used in a previous study [22]. For each pair of virus and host, we define the score *S*_*CRISP* *R*_(*v, b*) as the largest value of −log(E-value). If there is no match between a virus and host, a score of zero is assigned. We used `CRT1.2-CLI` [75] to find CRISPRs in all bacterial genomes, with parameters `-minNR 3 -minRL 20 -maxRL 50 -minSL 20 -maxSL 60 -searchWL 7`. All identified CRISPRs were merged to one file to construct a BLAST database using `makeblastdb` (`BLAST 2.6.0`). We then searched all viral genomes against the database using `blastn` with parameters `-evalue 1 -gapopen 10 -penalty -1 -gapextend 2 -word size 7 perc identity 90 -dust no -task blastn-short`.

With the CRISPR information, we modify the model of *π*_*ij*_ in Eq. (4) to

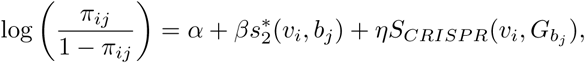

and our logistic regression model in Eq. (5) to

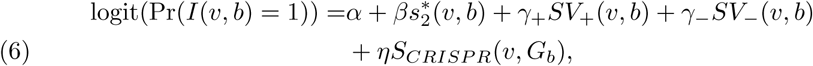

where *G*_*b*_ is the set of hosts that belong to the same genus as host *b*, and

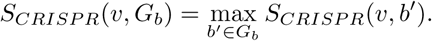

Due to the limited availability of CRISPR information in the training data, as shown in Fig. 2, we group hosts by genus for the CRISPR feature.

### The fraction of virus genome aligned to the host genome

Viruses and their hosts frequently exchange genetic materials and viruses play important roles in horizontal gene transfer. Therefore, similar regions in virus and host genomes can provide a strong evidence for linking a virus into its potential host. On the one hand, phages, especially those temperate phages, are able to integrate their own genomes to the hosts. On the other hand, phages can obtain genetic material from their hosts. If a genetic element brings an evolutionary advantage to the virus, the borrowed genetic segment will be preserved in the viral genome [22]. One example is cyanophages, phages that infect cyanobacteria. Many cyanophages acquire and express host photosystem genes that are thought to bolster photosynthetic energy during infection. [77].

Similar to the method in [22], we used `blastn` to find similarities between each pair of virus and host genomes. For each virus-host pair, their similarity, *S*_*blastn*_(*v, b*), is defined as the fraction of the virus genome that can be mapped to the host genome. Only matches with percent identity higher than 90% are used for prediction. Note that different parts of the virus genome can be matched to different positions on the host genome and all contribute to the coverage percentage. We used the same parameter setting as in [22] for our analysis. To generate blastn results a BLAST database was created for all bacterial genomes by makeblastdb (BLAST 2.6.0). We then searched all viral genomes against the database by `blastn` with parameters `-word size 11 -evalue 0.01 -reward 1 -penalty -2 -gapopen 0 -gapextend 0 perc identity 90`.

Finally, with the CRISPR feature and the alignment-based similarity, we have the following model:

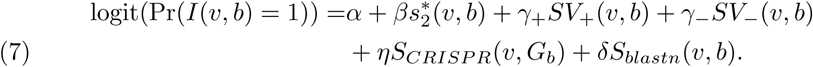

### Incorporation of WIsH score for predicting hosts of virus contigs

In many metagenomic studies, the whole genome of a virus may not be available. Instead, only parts of the virus genome referred as contigs that were assembled from shotgun reads are known. Several algorithms such as VirFinder and VirSorter etc. [78, 79, 80, 81, 82, 83] can be used to decide if the contigs come from virus genomes. Our objective is to predict the hosts for full virus genomes as well as viral contigs.

Galiez *et al.* [24] recently developed a program, WIsH, to predict the hosts of viral contigs and showed that WIsH outperforms 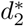 for predicting the hosts of viral contigs as short as 5 kb. WIsH trains a homogeneous Markov chain model for each host genome, and calculates the likelihood of a viral contig based on each Markov chain model. Instead of using 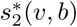 as a feature, we hereby replace it with the log-likelihood of viral contig *v* fitting to the Markov chain model of bacteria *b, S*_*WIsH*_ (*v, b*). WIsH [24] scores were computed using `WIsH 1.0` with the default parameters. Then the model for predicting the host *b* of viral contig *v* becomes,

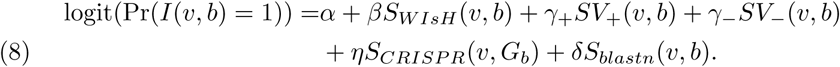

Note that both *SV*_+_(*v, b*) and *SV*_−_(*v, b*) are still computed by 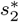, since WIsH is not able to depict the similarities between viral contigs.

### Model training and evaluation

Among the 1,427 viruses, we used the set of 352 viruses whose exact host genome sequences were known and the set of their corresponding 71 hosts as the positive training set. We randomly select 352 pairs of virus-host within the 352 viruses and 71 hosts as negative training data. To alleviate potential false negative interactions, we require that the selected host for each virus is not in the same phylum level as the true host. We then learned the model based on the training data for the various models. For real applications and the software, we set the coefficients by averaging over 100 times of the training procedure to reduce randomness.

It is possible that the selected 352 non-interacting pairs may contain some positive-yet-unknown interaction pairs, which may influence the training and test results. We recognized this possibility while assuming the fraction of such pairs is relatively low since the virus-host interaction is specific so that the overall fraction of virus-host interacting pairs among all the pairs is very small. The additional requirement that the host in a negative virus-host pair comes from a different phylum level further mitigates this potential problem.

The trained models are then used to predict the hosts of the remaining 1,075 viruses against 31,986 candidate prokaryotic hosts. For each virus, we estimate its probability of infecting any hosts, and the one with the highest probability was predicted as its host. For a taxonomic group 𝒮 at an upper taxonomic level containing a set of hosts, we define the prediction score between *v* and 𝒮 as the maximum probability between *v* and all hosts in 𝒮, that is

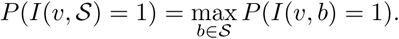

We predict the host group of the virus *v* by the one having the highest prediction score *P* (*I*(*v*, 𝒮) = 1). In case of ties, we first checked the number of hosts having the highest probability in each group and chose the one with the largest number of hosts having the highest probability. Further, if there were more than one taxon with the same number of bacteria having the highest probability, all taxa were reported.

We then compared the predicted host taxonomic groups with the true taxonomic group of every virus at several taxonomic levels: genus, family, order, class, and phylum. At a particular taxonomic level ℒ, let 𝒯_*v*_ be the set of predicted groups and *C*_ℒ_ (*v*) = *I*(*h*_*v*_, 𝒯_*v*_)*/*‖ 𝒯_*v*_‖, where *I*(*h*_*v*_, 𝒯_*v*_) = 1 if the true host of *v, h*_*v*_, belongs to the set of the predicted host groups 𝒯_*v*_, and *I*(*h*_*v*_, 𝒯_*v*_) = 0, otherwise. The prediction accuracy for a certain taxonomic level is defined as

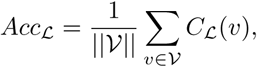

where 𝒱 is the set of viruses for prediction.

### Clustering of viral contigs

To examine the relatedness of viral contigs for novel host predictions, proteins encoded on viral contigs were predicted by `Prodigal 2.6.3` (with default parameters). `BLASTp 2.6.0` was then used to search for similar proteins shared between viral contigs. The percentage of genes shared between two contigs were defined as the number of pairs of homologous proteins between the two contigs divided by the average number of proteins of the two contigs.

### Consideration of virus-host co-abundance in host prediction

In order to investigate whether co-abundance can help the prediction of virus-host interactions, we incorporated this feature to the model in a smaller data set to evaluate its contribution. The data set included a subset of 2,695 prokaryotic reference genomes and 1,403 viruses (see below). A total of 148 stool metagenomic samples from the Human Microbiome Project (HMP) [84] and 103 metagenomes from the Tara Ocean (filter size 0.22 to 3 *µ*m) [85] were collected. We used `centrifuge` [81] (`centrifuge-1.0.3-beta`) to compute the abundance of virus and bacteria genomes in each of the metagenomes, resulting in an abundance profile of 251-dimensional vector for every virus and host genome. The co-abundance feature *S*_*co*−*abundance*_(*v, b*) was defined by the Pearson correlation between the abundance profiles for the pair of virus and bacterium. We then modified the integrated model to

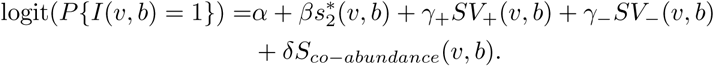

We compared the performance of this model with that of the model in Eq. (5). Both models were trained based on a subset of 308 viruses and 50 hosts, including 308 pairs of true interacting pairs (instead of 352 known interacting pairs used in the main text, because the hosts of the other 44 pairs are not in the 2,695 host genomes) and 308 randomly chosen negative pairs. After both models were trained, we predicted the hosts of 1,095 viruses. The results are shown in Additional File 14. The co-abundance feature itself had weak prediction ability and adding it to the model did not help prediction. Therefore, we did not consider it as a feature in the final model presented in the main text.

### Software

We developed a computational tool, VirHostMatcher-Net, implementing our network-based integrated method for virus-host predictions. The software is publicly available at https://github.com/WeiliWw/VirHostMatcher-Net. The tool supports parallel computing and has the option of choosing the type of query viruses (complete genomes or contigs). It also provides the option of specifying a customized subset of candidate hosts for prediction. The tool provides informative outputs including all the feature scores of the query viruses against all candidate hosts, and a summarized table listing top predictions for each virus with their feature scores, score percentiles, and accuracy. The score percentile of a virus-host pair is defined as the percentile of this score among all scores between that virus and all the candidate hosts. A large percentile suggests high relevance of the feature score. The percentile of *SV*_−_, the only feature with a negative coefficient, is reversed to be consistent with other feature score percentiles. The percentile information helps to better understand how relevant each feature score is for a particular prediction. We also provide “accuracy” that gives the fraction of correct predictions when virus-host pairs with prediction scores above the particular threshold are declared as interacting.

## Data and code availability

Accession numbers for viral contigs in the real data studies can be found in the section of Data sets. All other relevant data for training and testing the model and code are available at https://github.com/WeiliWw/VirHostMatcher-Net.

## Author’s contributions

J.R., F.S., and N.A.A. conceived the study and designed the method. W.W. and J.R. developed the algorithms and implemented the data analysis. N.A.A. led the analysis of the biological data sets. W.W., J.I-E, F.S., and N.A.A. wrote the manuscript. W.W. developed the software and K.T. helped in improving the performance. E.D., J.I-E., N.A.A., J.A.F., J.B., and J.I-E. helped in discussing and interpreting the results, and revised the manuscript critically. F.S. and N.A.A finalized the paper. All authors read and approved the final manuscript.

## Acknowledgements

We thank Dr. Michael S. Waterman at the University of Southern California (USC) for helpful discussions. Dr. Yang Lu and Dr. Mengge Zhang at USC participated in the discussions and some of the calculations at the beginning of the project.

## Additional Files

**Additional file 1: Additional file 1.pdf.** Supplemental figure in PDF format. ROC curves for predicting virus-host interactions using 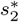 and WIsH on 352 positive and negative virus-host pairs.

**Additional file 2: Additional file 2.pdf.** Supplemental figure in PDF format. Improvement in host prediction by thresholding on the prediction score for viral contigs of different lengths across different families of Caudoviruses.

**Additional file 3: Additional file 3.csv.** Supplemental table in CSV format. The prediction of ΦcrAss001 from Shkoporov *et al.* The table lists the highest scores for 22 host species tested in Shkoporov *et al.*.

**Additional file 4: Additional file 4.csv.** Supplemental table in CSV format. The predicted hosts of 249 crAss-like phages from Guerin *et al.*. The column “cluster” lists the original genus annotated in Guerin *et al.*.

**Additional file 5: Additional file 5.pdf.** Supplemental figure in PDF format. The recruitment plot of *Akkermansia muciniphila* sequences to the crAssphage genome.

**Additional file 6: Additional file 6.csv.** Supplemental table in CSV format. The predictions of the 1,238 EVGs from Nishimura *et al.* with prediction scores ≥0.78 by our integrated model. In the table, the five columns ending with “val” list the corresponding feature scores; the five columns starting with “acc” list the empirical accuracies at different taxonomic levels; the five columns ending with “pct” list the score percentiles of the scores for the corresponding features between the virus and all hosts. Note when the feature is not available for all the hosts, the corresponding percentile will leave blank. The last column “overlap” represents if the virus was previously annotated in Nishimura *et al.*. Columns with “Nishimura.” report the host predictions made by Nishimura *et al.*.

**Additional file 7: Additional file 7.csv.** Supplemental table in CSV format. The predicted hosts of 9,625 human associated viral contigs from Paez-Espino et al. with scores ≥0.78 by the integrated model. The hosts of all of these viral contigs were not predicted previously and were predicted exclusively by our method.

**Additional file 8: Additional file 8.csv.** Supplemental table in CSV format. The predicted hosts of 31,041 marine viral contigs from Paez-Espino et al. with scores ≥0.78 by the integrated model. The hosts of all of these viral contigs were not predicted previously and were predicted exclusively by our method.

**Additional file 9: Additional file 9.csv.** Supplemental table in CSV format. The predicted hosts of 20,643 viral contigs in other environments from Paez-Espino et al. with scores ≥0.78 by the integrated model. The hosts of all the viral contigs were not annotated previously and were predicted exclusively by our method.

**Additional file 10: Additional file 10.csv.** The predicted hosts of the selected group of 152 human associated viral contigs from Paez-Espino et al. with scores ≥0.78 by the integrated model. The hosts of all of these viral contigs were not predicted previously and predicted exclusively by our method.

**Additional file 11: Additional file 11.pdf.** Supplemental figure in PDF format. Relatedness of newly discovered 152 viral contigs in human associated metagenomes based on shared gene content.

**Additional file 12: Additional file 12.csv.** Supplemental table in CSV format. The predicted hosts of the selected group of 350 marine viral contigs from Paez-Espino et al. with scores ≥0.78 by the integrated model. The hosts of all of these viral contigs were not predicted previously and were predicted exclusively by our method.

**Additional file 13: Additional file 13.pdf.** Supplemental figure in PDF format. Shared gene relatedness of 350 newly discovered viral contigs and 31 previously isolated viruses that infect *Cellulophaga*.

**Additional file 14: Additional file 14.pdf.** Supplemental figure in PDF format. Investigation of prediction accuracy using virus and host co-abundance.

**Additional file 15: Additional file 15.pdf.** Supplemental figure in PDF format. The effect of prediction accuracies when the hosts in the true genus level are excluded.

