## Supplementary figures and images for "A network-based integrated framework for predicting virus-host interactions"

### Additional_file_1.pdf

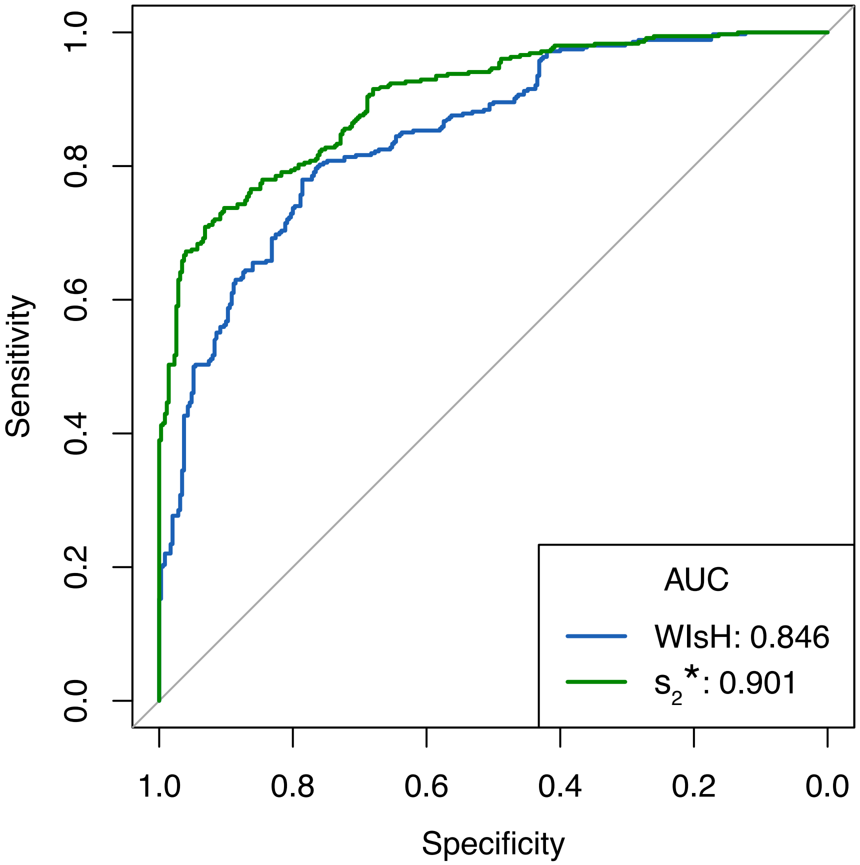

### Additional_file_2.pdf

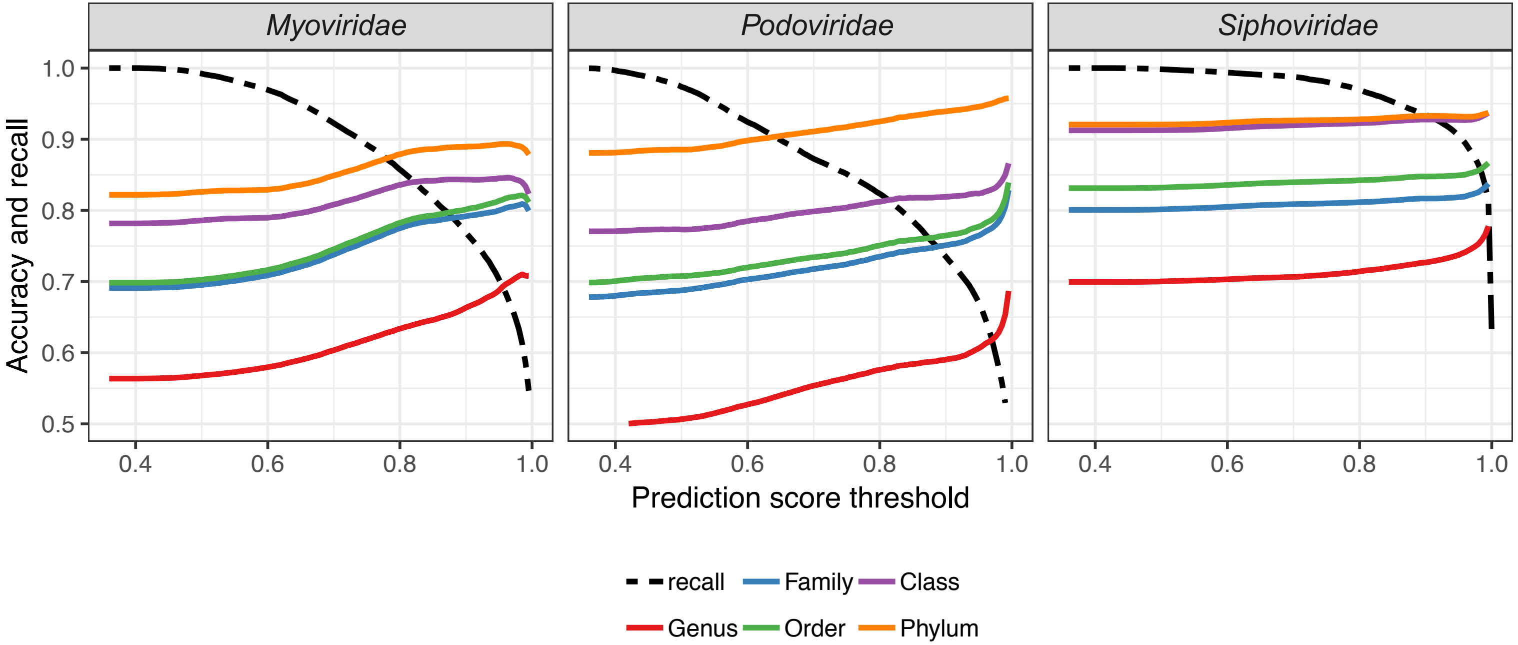

### Additional_file_5.pdf

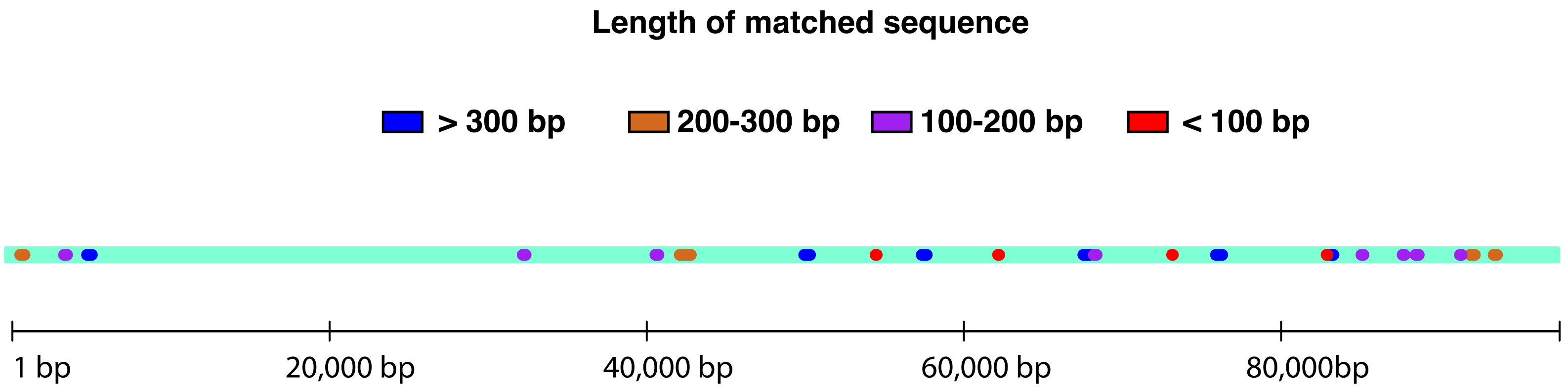

### Additional_file_11.pdf

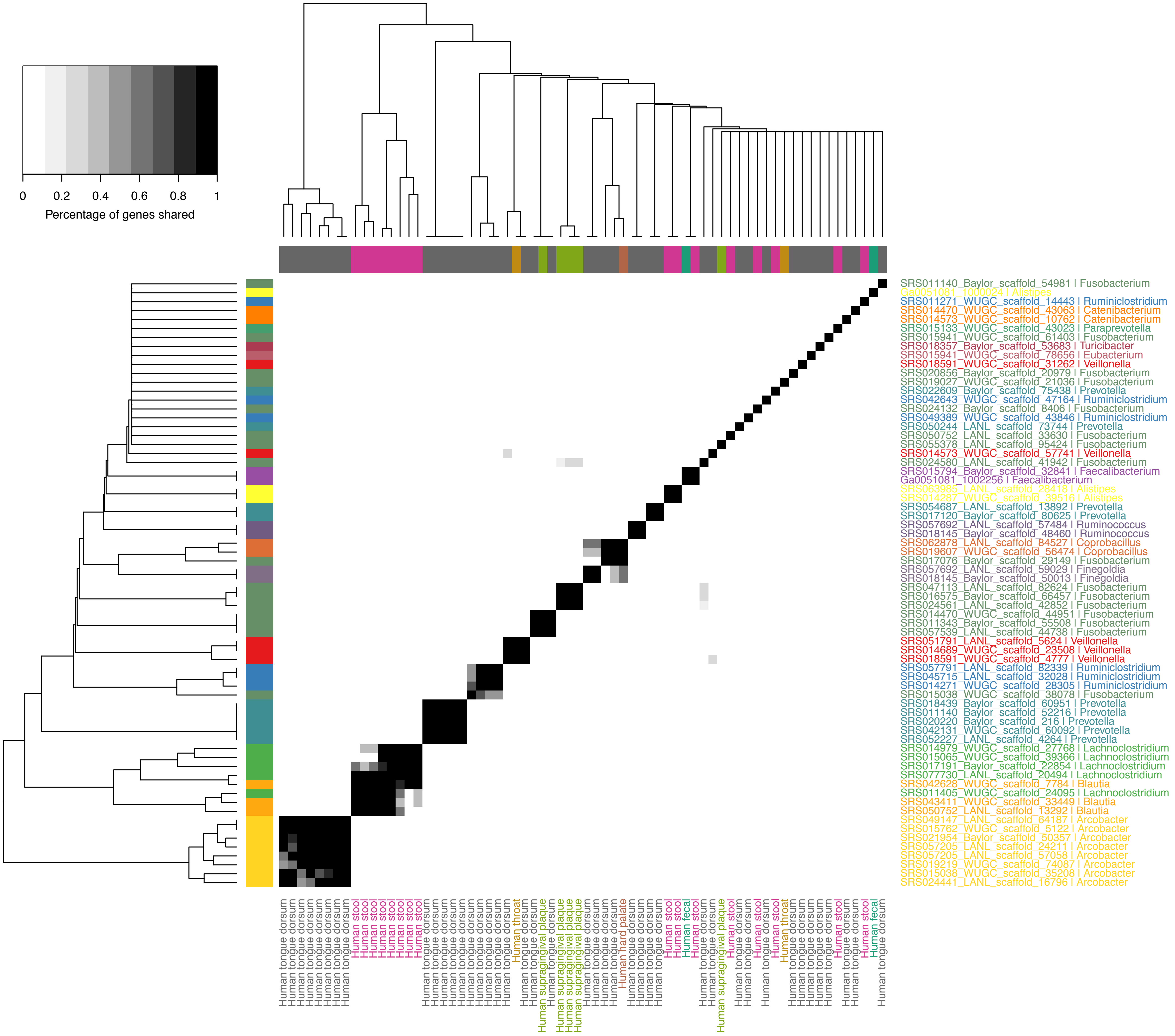

### Additional_file_13.pdf

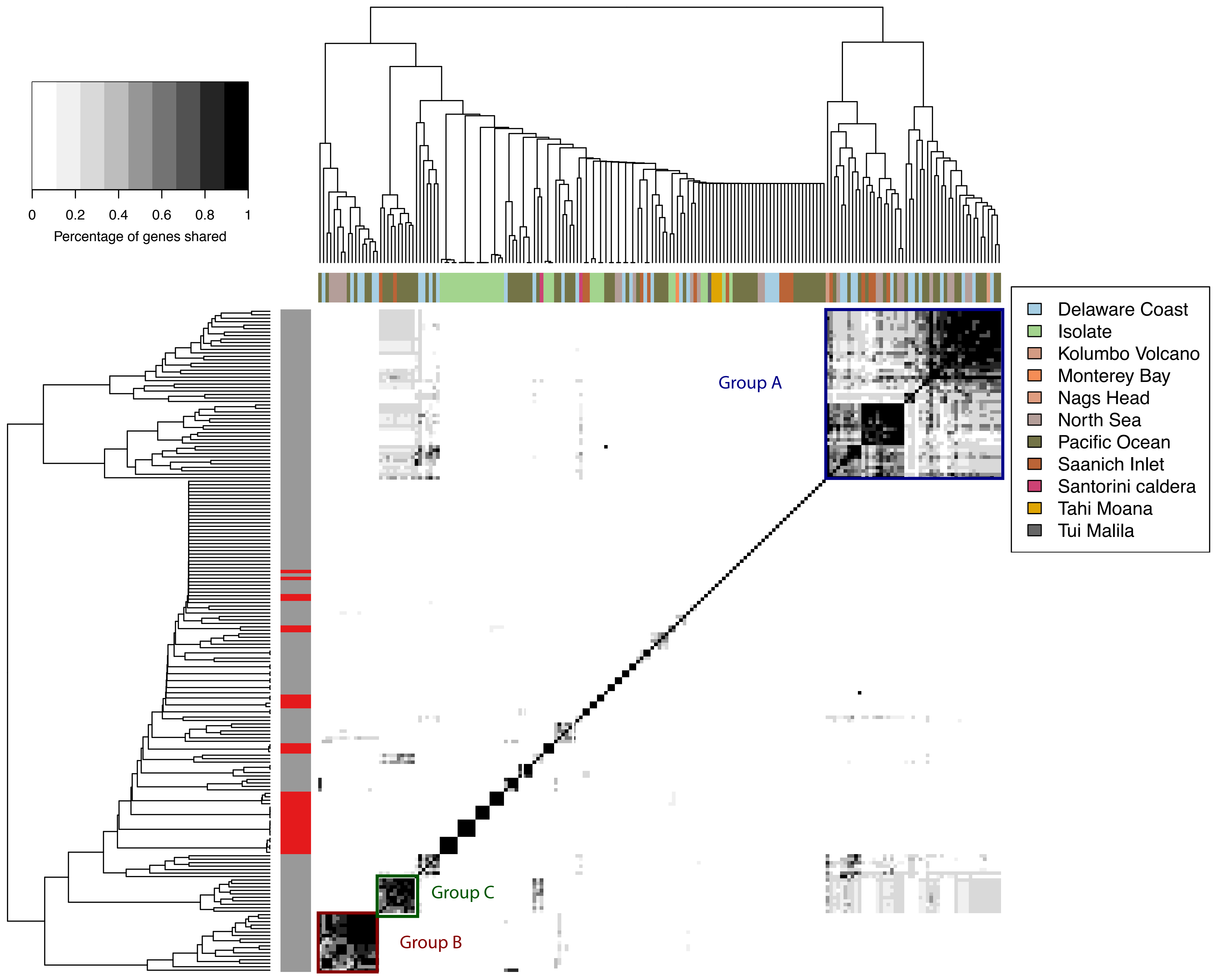

### Additional_file_14.pdf

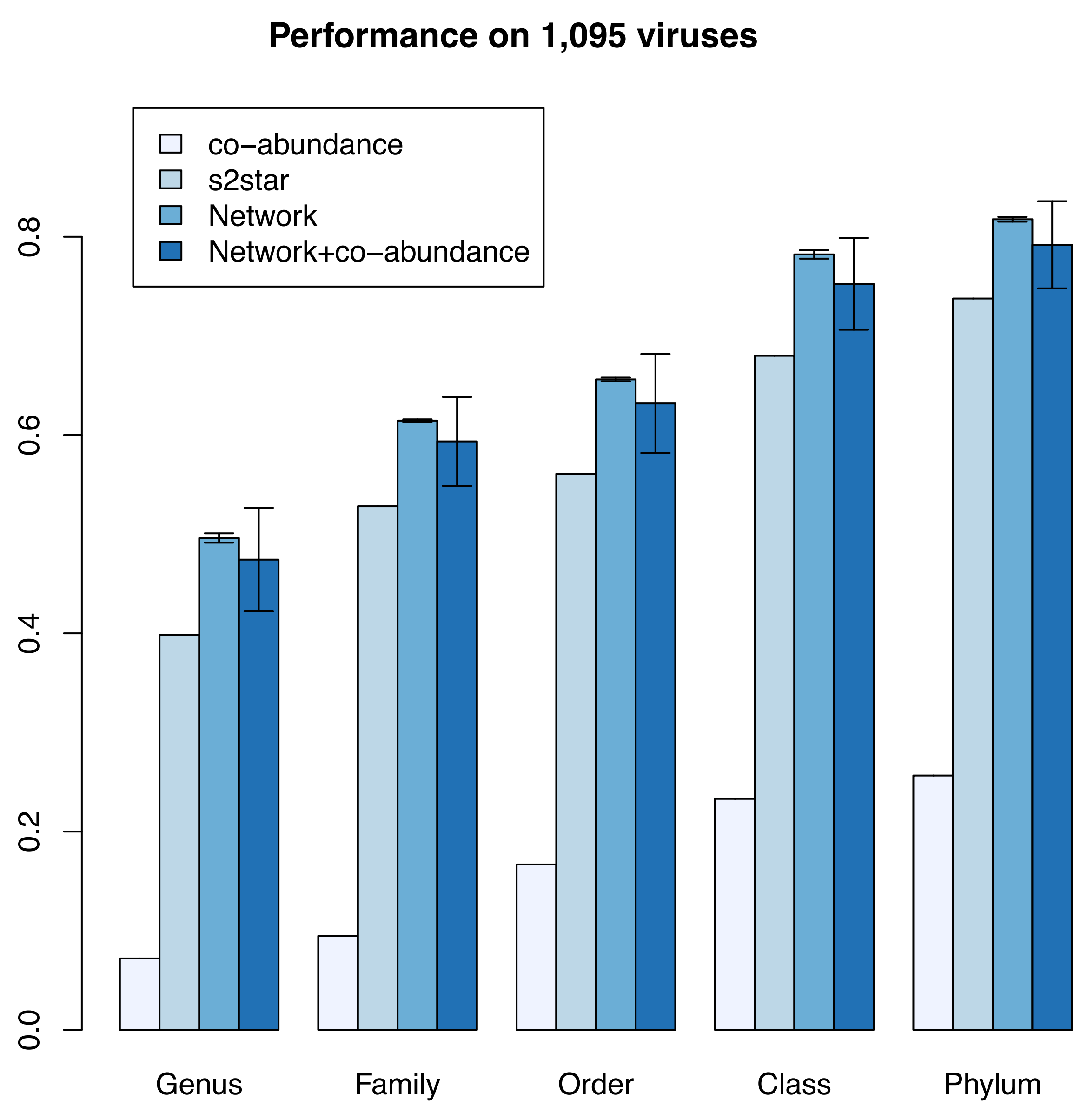

### Additional_file_15.pdf

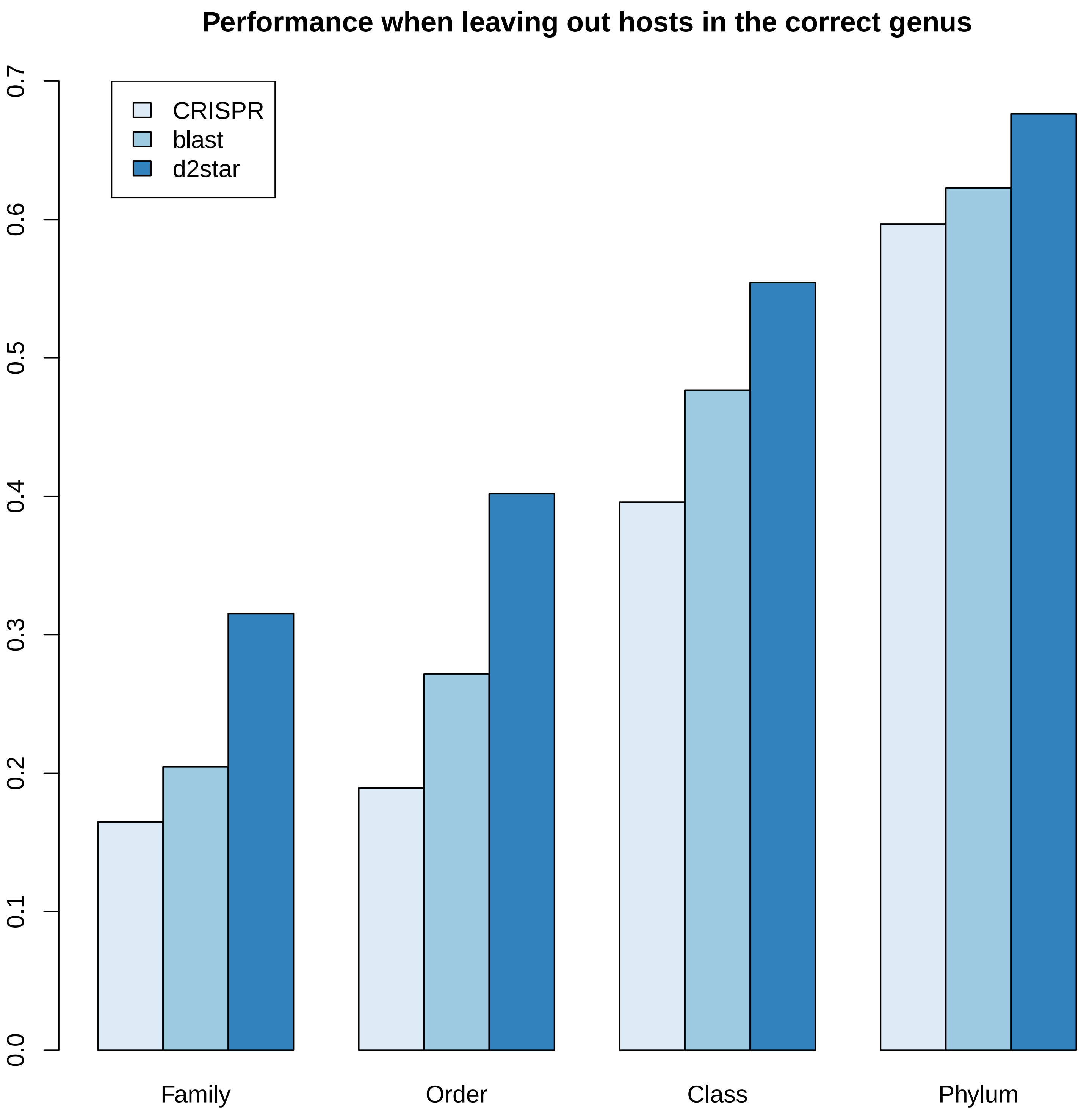
